# Physical properties of the cytoplasm modulate the rates of microtubule polymerization and depolymerization

**DOI:** 10.1101/2020.10.27.352716

**Authors:** Arthur T. Molines, Joël Lemière, Morgan Gazzola, Emilie I. Steinmark, Claire H. Edrington, Chieh-Ting (Jimmy) Hsu, Klaus Suhling, Gohta Goshima, Liam J. Holt, Manuel Thery, Gary. J. Brouhard, Fred Chang

## Abstract

The cytoplasm is a crowded, visco-elastic environment whose physical properties change according to physiological or developmental states. How the physical properties of the cytoplasm impact cellular functions *in vivo* remain poorly understood. Here, we probed the effects of cytoplasmic concentration on microtubules by applying osmotic shifts to fission yeast, moss, and mammalian cells. We show that both the rates of microtubule polymerization and depolymerization scale linearly and inversely with cytoplasmic concentration; an increase in cytoplasmic concentration decreases the rates of microtubule polymerization and depolymerization proportionally, while a decrease in cytoplasmic concentration leads to the opposite. Numerous lines of evidence indicate that these effects are due to changes in cytoplasmic viscosity rather than cellular stress responses or macromolecular crowding *per se*. We reconstituted these effects on microtubules *in vitro* by tuning viscosity. Our findings indicate that, even in normal conditions, the viscosity of cytoplasm modulates the reactions underlying microtubule dynamic behaviors.

## Introduction

Cytoplasm is composed of 100-300 mg/ml of macromolecules (proteins, nucleic acids, lipids, etc.), which occupy 10-40% of the total cellular volume (Milo and Phillips, 2015; Neurohr and Amon, 2020). These components range in size from small globular proteins to extended networks of organelles and cytoskeletal polymers. Ribosomes alone occupy ∼20% of that volume (Delarue et al., 2018). Biophysical studies revealed that these constituents form a porous, visco-elastic material that allows diffusion of small molecules but impedes movement of larger particles (Luby-Phelps et al., 1986; Moeendarbary et al., 2013; Xiang et al., 2020), and molecular simulations show a high density of macromolecules jostling and colliding from diffusive motion. (Mcguffee and Elcock, 2010; Yu et al., 2016).

In contrast to the density of inert materials, the density of the cytoplasm is regulated as part of cell physiology. Indeed, cytoplasmic density varies during the cell cycle, among different cell types, as a result of aging, as a response to nutritional stress, and as a response to disease (Neurohr and Amon, 2020). These density changes likely affect macromolecule concentrations and the physical properties of the cytoplasm such as its degree of crowding and/or its viscosity, which in turn will impact a broad range of cellular processes, such as protein-protein associations, phase transitions, and enzymatic fluxes. It was also recently proposed that cells can tune the viscosity of their cytoplasm to regulate diffusion-dependent processes in response to temperature (Persson et al., 2020). Thus, it is critical to understand how cellular reactions are affected by the physical properties of cytoplasm.

The most prominent physical property of cytoplasm is macromolecular crowding. There are several conceptual models used to describe the influence of macromolecular crowding on biochemical reactions. Minton and colleagues have argued that bulky macromolecules “exclude volume”, a steric effect that increases the thermodynamic activity of other proteins (Minton, 2006; Shahid et al., 2017). Consistent with this idea, bulky macromolecules often accelerate biochemical reactions *in vitro*. Additionally, because macromolecules are rarely inert, they interact with proteins via short-term, non-specific hydrophobic interactions that can affect the rates and equilibria of reactions (Mcguffee and Elcock, 2010). Theoretical models of macromolecular crowding can explain how crowding impedes diffusion, produces entropic forces that draw molecules together, promotes phase separation, and produces osmotic pressure within cells (Mitchison, 2019; R.John, 2001; Shahid et al., 2017). But the models of macromolecular crowding do not always make the same predictions. For example, Mitchison used the concept of colloidal osmotic pressure to argue that cytoplasm is functionally dilute, such that a reaction like microtubule (MT) polymerization would be unaffected by steric effects *in vivo* (Mitchison, 2019). To distinguish between these concepts, what is needed are experiments that perturb the physical properties of cytoplasm and measure the rates of cellular reactions *in vivo*.

The dynamic behavior of MTs represents an attractive case with which to probe the effects of the cytoplasm on defined biochemical reactions *in vivo*. First, the polymerization and depolymerization of single MTs are reactions that can be quantitatively measured in living cells and *in vitro* using microscopy. Second, the effects of macromolecular crowding on MTs *in vitro* are known: MTs grow significantly faster in the presence of bulky crowders but significantly slower in the presence of small viscous agents like glycerol (Wieczorek et al., 2013). These *in vitro* measurements can be compared to *in vivo* measurements when considering mechanisms. Third, at 100 kDa in mass and 8 nm in length, the tubulin dimer represents a size range typical for soluble proteins and enzymes, while the MT diameter at around 25 nm is similar to large macromolecular complexes such as the ribosome. Finally, MT polymerization depends on tubulin concentration while MT depolymerization does not (Fygenson et al., 1994; Walker et al., 1988). Thus, changes in tubulin concentration (e.g., because of changes in cytoplasmic concentration) should impact polymerization alone, which is a testable prediction of some models. Taken together, MTs are well-suited to probe what properties of cytoplasm have the strongest impact on cellular reactions, and more generally, to inform biophysical models describing the physical properties of cytoplasm.

Fission yeast is an excellent model organism with which to study the physical regulation of MT dynamics *in vivo*. We can readily image the interphase MT bundles and measure the dynamic behavior of individual MTs with precision (Höög et al., 2007; Loiodice et al., 2019; Sawin and Tran, 2006). Importantly, we can readily manipulate the properties of their cytoplasm using osmotic shifts, which create robust and well characterized changes in cellular volume and cytoplasmic concentration (Atilgan et al., 2015; Knapp et al., 2019).

Here, we study the effects of the physical properties of the cytoplasm on MTs dynamics by using acute osmotic shifts to vary cytoplasmic concentration. We show that hyperosmotic shifts, which increase cytoplasmic concentration, lead to dampening and “freezing” of MT polymerization and depolymerization. Conversely, hypoosmotic shifts, which decrease cytoplasmic concentration, lead to increased rates of MT polymerization and depolymerization. The observed proportionate changes to MT rates, which were independent of the osmotic stress response and key MT regulators, correlated with global changes in cytoplasmic physical properties and were recapitulated *in vitro* through modulation of viscosity. These findings demonstrate that cytoplasm modulates MT dynamics through viscous effects even at normal concentrations of cytoplasm.

## Results

### Cytoplasmic concentration tunes microtubule dynamics

The density of cellular components can be experimentally manipulated by varying the osmotic environment using osmotic agents such as sorbitol (Knapp et al., 2019). We manipulated live fission yeast cells by adding sorbitol to the growth medium, which led to an acute decrease in cell volume in a dose-dependent, reversible manner (Fig. 1 A and B) (Sup. Fig. 1 and Sup. Fig. 2) (Atilgan et al., 2015; Knapp et al., 2019). For instance, volume decreased by up to 50% with 1.5 M sorbitol added to rich media, without loss in cell viability (Sup. Fig. 1). This volume decrease presumably occurs through water loss. We confirmed the volume decreases by measuring increases in fluorescence intensity of GFP-labeled tubulin and a ribosomal protein Rps802 (Sup. Fig. 1).

**Figure 1:**
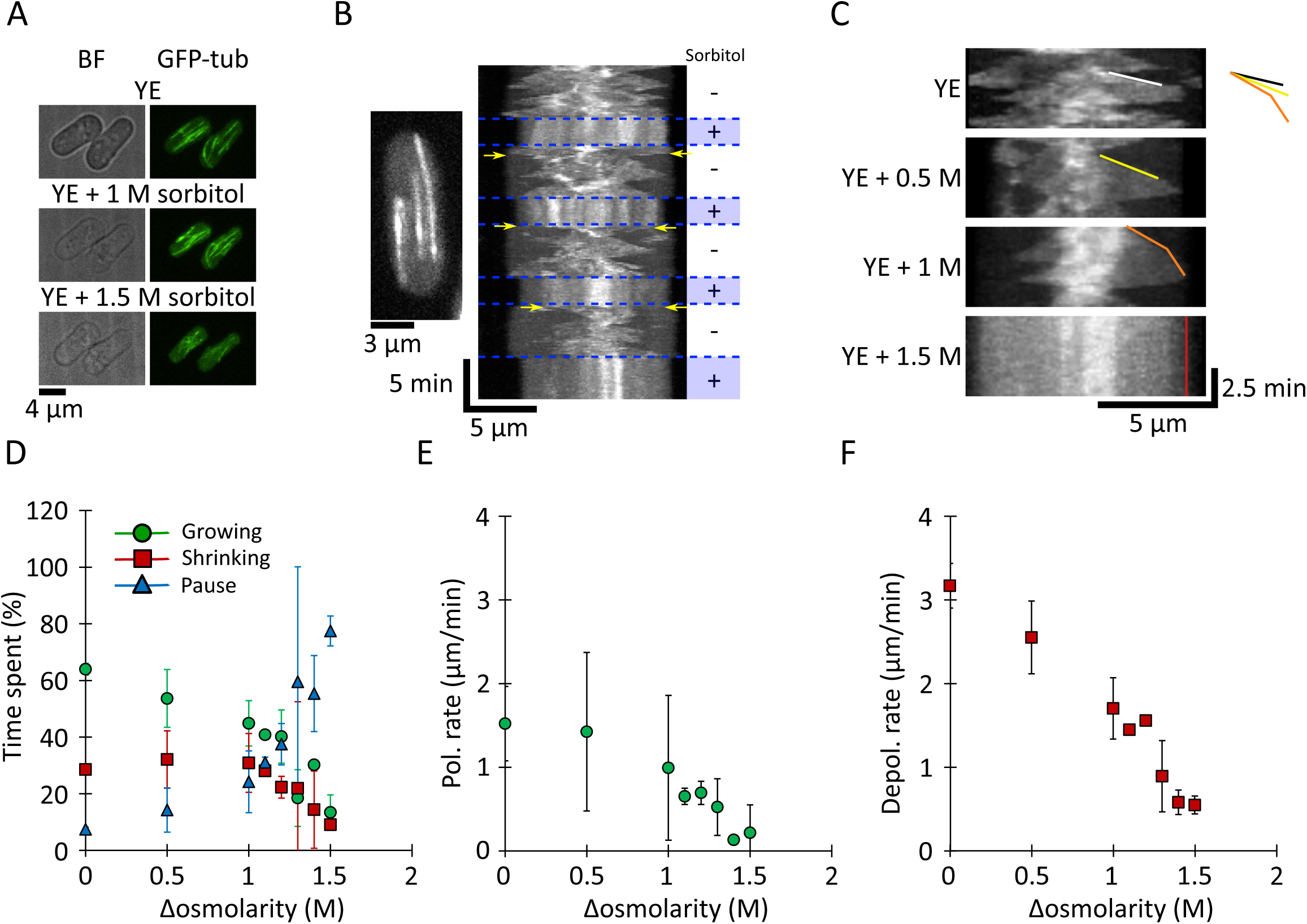
**MT growth and shrinkage rates decrease upon hyperosmotic shock in yeast cells.** (A) Brightfield (BF; left) and fluorescence (right) images of interphase MT bundles in two fission yeast cells expressing GFP-tubulin (GFP-tub) upon sequential treatment with YE (medium alone), YE + 1 M sorbitol, and YE + 1.5 M sorbitol. (B) MT dynamics in a cell treated with oscillations of YE (5 min) and YE + 1.5 M sorbitol (3 min). In the kymograph, this representative cell is expressing GFP-tubulin; the cell image has been collapsed onto a line. MTs exhibit decreased dynamics acutely in 1.5 M sorbitol. Upon sorbitol washout, MTs first depolymerize (yellow arrows) and then resume dynamic behaviors. (C) Kymographs of MTs in yeast cells at the indicated sorbitol concentrations. Lines highlight tracks of single growing MT plus-ends. Colored lines highlight growth events. (D) Percentage of time that interphase MTs spent growing (green circles), shrinking (red squares), or paused (blue triangles) in YE and after hyperosmotic shocks (AVG +/- standard deviation). Cell and MT values of n are as in (E). (E) MT polymerization rates (green circles) in yeast cells treated acutely with the indicated sorbitol concentrations. Values are AVG +/- standard deviation. Data come from (left to right) n = 58 / 51 / 32 / 27 / 38 / 23 / 15 / 26 cells and 118 / 99 / 60 / 65 / 72 / 34 / 22 / 44 MT, respectively, from at least 2 experiments. (F) MT depolymerization (red squares) rates in yeast cells treated acutely with the indicated sorbitol concentrations. Values are AVG +/- standard deviation. Cell and MT values of n are as in (E).

Having validated this approach to alter cytoplasmic concentration, we applied it to cells expressing GFP-tubulin to monitor MT dynamics. Time-lapse imaging of untreated cells showed that interphase MTs were characteristically dynamic, polymerizing and depolymerizing, with little time in “pause” (Tran et al., 2001). In contrast, hyperosmotic shifts caused significant changes in the dynamic behaviors of interphase MTs, as noted previously (Robertson and Hagan, 2008; Tatebe et al., 2005). We found that in acute response to hyperosmotic shifts, the interphase MT cytoskeleton appeared to “freeze” (Fig. 1B and C), especially at high sorbitol concentrations (1.5 M) (Fig. 1 B, C and D). In general, MTs were “paused” at various lengths and exhibited little or no polymerization or depolymerization. To determine whether the effects of the hyperosmotic shifts were reversible, we cycled the concentration of sorbitol from 0 to 1.5 M in 5 min intervals. Upon each hyperosmotic shift to 1.5 M sorbitol, most of the MTs “froze” within 30 sec (Fig. 1 B and C). Upon each shift back to sorbitol-free medium, all interphase MTs promptly went into catastrophe, shrank toward the middle of the cell, and then regrew, such that the normal interphase array was restored in a few minutes (< 5 min) (Fig. 1 B and Sup. Fig. 2). The prompt catastrophes suggest that the GTP cap is hydrolyzed when MTs are in the “frozen” state. This cycle of MT “freezing”, and resumption of dynamics could be induced repeatedly (Fig. 1 B and Sup. Fig. 2), demonstrating that the effects of hyperosmotic shifts were rapid and reversible.

The effects on MT dynamics were dependent on the sorbitol concentration. We detected a progressive increase in the time that MTs spent in a pause state (Fig. 1 D). Without sorbitol MTs spent 7 +/- 1 % of the time in pause, while at 1 and 1.5 M sorbitol they spent 24 +/- 11 % and 77 +/- 5 % of the time in pause respectively (Fig. 1 D). Of the MTs that continued to be dynamic, their rates of polymerization and depolymerization decreased in a sorbitol dose-dependent manner (Fig. 1 C, E, and F, and Sup. Fig. 3). Importantly, sorbitol’s effects on MT polymerization and depolymerization were equivalent in magnitude, a point to which we will return later. For instance, at 1.5 M sorbitol, polymerization and depolymerization rates decreased by -79 +/- 2 % and 80 +/- 1 %, respectively (Fig. 1 E and F). Because both polymerization and depolymerization rates were affected, we can rule out a mechanism based on changes in the concentration of tubulin, which should affect polymerization only. Treatment with high sorbitol also made MTs resistant to depolymerization by cold temperature (Sup. Fig. 4), further indicating that these MTs were in a highly stabilized state.

We next asked whether hypoosmotic shifts, which dilute the cytoplasm by causing influx of water, yield opposite effects on MT dynamics. Our initial attempts to swell intact fission yeast cells with hypoosmotic shifts were not successful, perhaps because the cell wall limited the swelling. However, the cell wall can be removed enzymatically to create protoplasts, which swelled substantially in response to hypoosmotic shifts without lysing (Fig. 2 A) (Lemière and Berro, 2018). In order to establish a control condition for protoplasts, we determined that protoplasts produced in 0.4 M sorbitol in rich media had the same average volume as intact fission yeast cells. Shifting the protoplasts to hypoosmotic and hyperosmotic conditions (relative to 0.4 M sorbitol) led to predictable changes in cell volume over a ∼2.5-fold range (Fig. 2 A). MTs were readily imaged across all volumes (Fig. 2B). In hypoosmotic shifts, MT polymerization and depolymerization rates both increased by an equivalent magnitude (Fig. 2C to 2F). For example, when cell volume swelled by 56 +/- 1 % over isotonic conditions, polymerization rates increased by 64 +/- 3 % and depolymerization rates increased by 42 +/- 1 % relative to control (Fig. 2 E and F). Conversely, hyperosmotic shifts in protoplasts decreased the rates of MT polymerization and depolymerization, also with equivalent magnitudes, similar to what was observed in intact fission yeast cells (Fig. 2 C to F). This result implies that the properties of the cytoplasm dampen the rates of MT polymerization and depolymerization.

**Figure 2:**
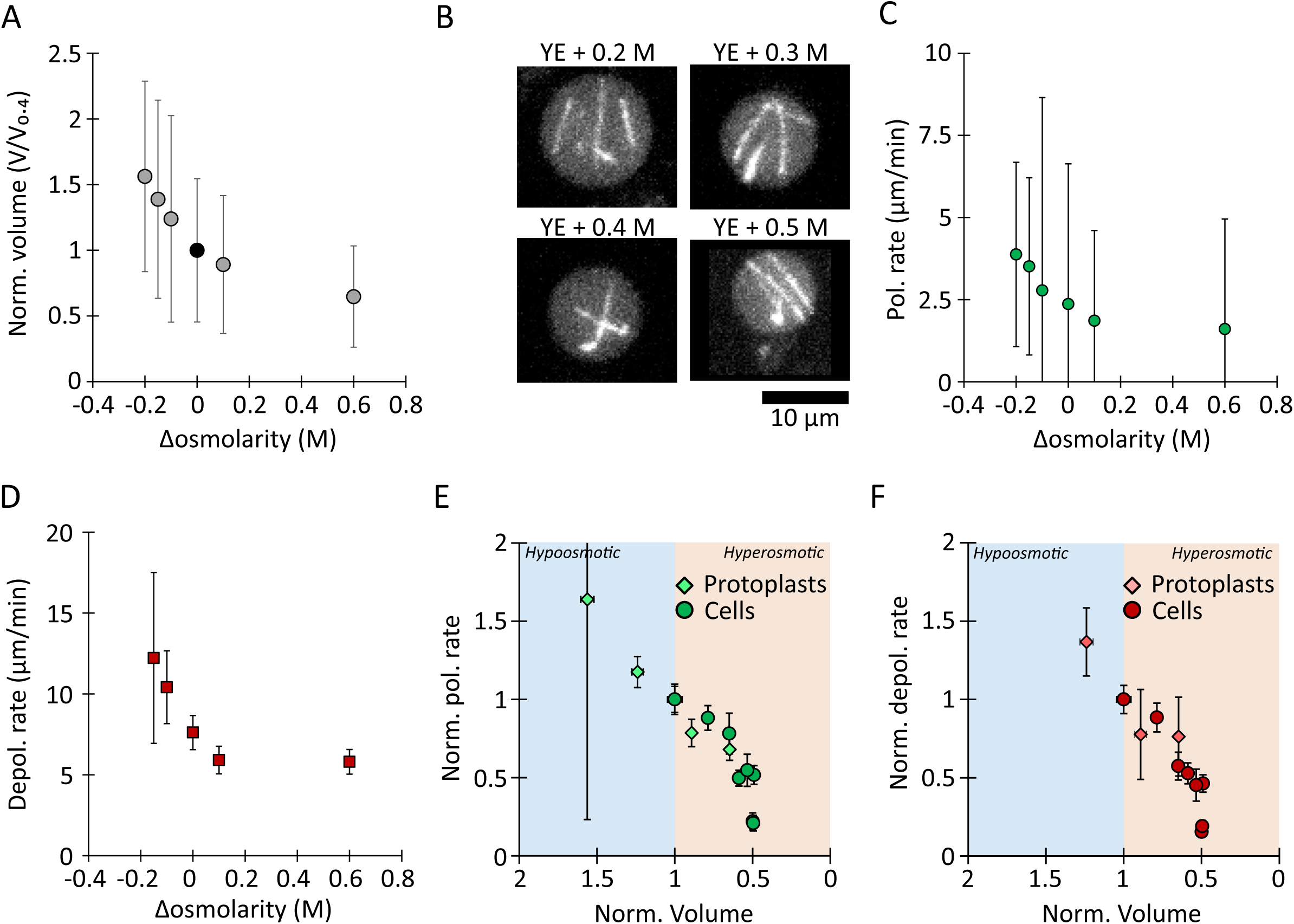
**MT growth and shrinkage rates scale with cell volume in yeast protoplasts.** (A) Normalized protoplast volume. Data were normalized to osmolarity conditions of YE + 0.4 M sorbitol, in which the average volume distribution of protoplasts matched that of intact cells (AVG +/- standard deviation). Data come from at least two experiments; (left to right) n = 18 / 45 / 109 / 446 / 200 / 440 protoplasts. (B) Representative images of GFP-labelled MTs in protoplasts in medium supplemented with the indicated sorbitol concentrations. (C) MT polymerization (Pol.) and (**D**) depolymerization (Depol.) rates in yeasts protoplasts in medium supplemented with the indicated sorbitol concentrations. Values are AVG +/- standard deviation. Left to right, n = 10 / 13 / 64 / 29 / 28 / 25 polymerization events and n = 7 / 57 / 25 / 13 / 12 depolymerization events from three experiments. (**E**) MT polymerization rate and (**F**) depolymerization rates, normalized to the isotonic condition for yeast cells (circles) and yeast protoplasts (diamonds), as a function of the normalized volume (see Methods). Both rates are increase in hypo-tonic conditions (blue shading) and decrease in hyper-tonic conditions (orange shading).

To evaluate whether cytoplasmic concentration sets the rates of dynamic instability in other cell types, we performed similar osmotic shifts with the moss *Physcomitrium (Physcomitrella) patens* and mammalian Ptk2 cells. In both cases, we observed lower MT polymerization and depolymerization rates after hyperosmotic shifts, similar to what we observed in fission yeast (Fig. 3 and Sup. Fig. 5). In Ptk2 cells (Fig. 3) treated with DMEM + 0.25 M sorbitol, MT polymerization rate decreased from 4.9 +/- 1.5 μm/min to 2.4 +/- 0.8 μm/min (-48%) while depolymerization rate decreased from 12 +/- 3.4 μm/min to 6 +/- 2.5 μm/min (-50%) (mean +/- standard deviation). In moss cells (Sup. Fig. 5) treated with BCD + 0.5 M sorbitol MT polymerization rate decreased from 5 +/- 1 μm/min to 4 +/- 0.7 μm/min (-20%) while depolymerization rate decreased from 30 +/- 19 μm/min to 22 +/- 15 μm/min (-28%) (mean +/- standard deviation). The similar effects of osmotic shifts on MT in fungal, plant and mammalian cells suggest that they arise from a conserved mechanism.

**Figure 3:**
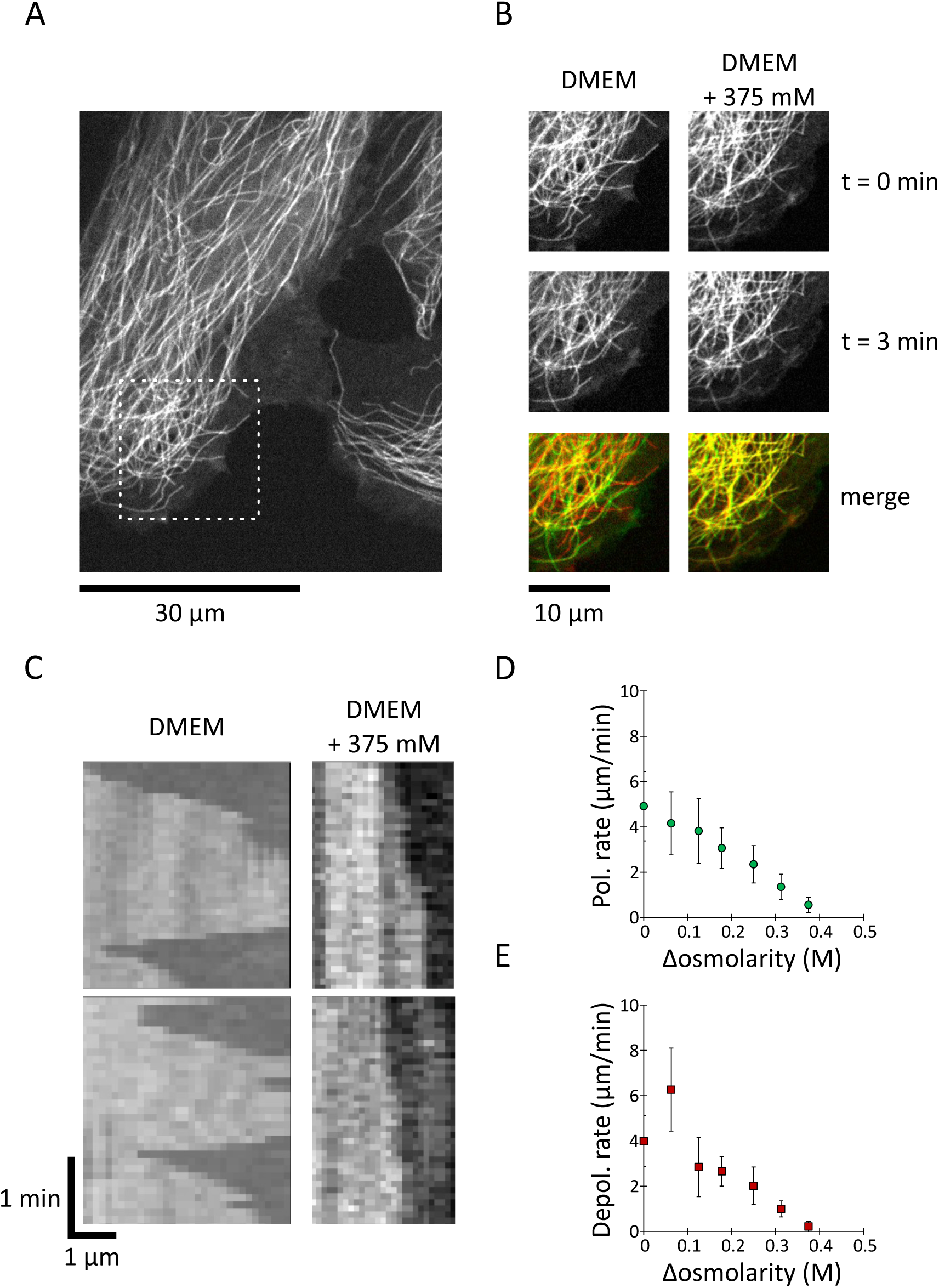
**MT growth and shrinkage rates decrease upon hyperosmotic shock in mammalian cells.** (A) Representative images of Ptk2 cells stably expressing GFP-labelled tubulin. Dashed-boxed ROI is zoomed-in in (**B**). (B) Snapshots from the cell in (**A**) showing MT dynamics before and after an osmotic shock with DMEM/F12 media containing 375 mM of sorbitol. (C) Kymographs of MTs in Ptk2 cells at the indicated sorbitol concentrations. (D) MT polymerization rates (green circles) in Ptk2 cells treated acutely with the indicated sorbitol concentrations. Values are AVG +/- standard deviation. Data come from (left to right) n = 39 / 58 / 59 / 48 / 57 / 61 / 40 MT, from at least 3 cells, from at least 2 experiments. (E) MT depolymerization rates (red squares) in Ptk2 cells treated acutely with the indicated sorbitol concentrations. Values are AVG +/- standard deviation. Data come from (left to right) n = 29 / 47 / 49 / 48 / 47 / 41 / 24 MT, from et least 3 cells, from at least 2 experiments.

Normalization of the MT polymerization and depolymerization rates to the isotonic conditions for both yeast cells and protoplasts revealed how changes in the concentration of the cytoplasm, above and below normal levels, caused similar, linear responses for MT polymerization and depolymerization (Fig. 2 E and F). This response suggests that the property of the cytoplasm that is changed by osmotic shocks affects both MT polymerization and depolymerization in a similar manner. Interestingly, MT dynamics decreased when the cytoplasm concentration and tubulin concentration increased (Fig. 1 and 2), opposite to what would be expected if crowding or tubulin concentration were responsible for MT dynamics as MT polymerization increases with either of these parameters (Wieczorek et al., 2013).

### Effect of osmotic shifts on MT dynamics is independent of stress response and regulatory proteins at MT plus ends

Cytoplasmic concentration could influence MTs directly, through its physical properties, or indirectly, e.g., through osmotic stress response pathways. We next investigated several plausible mechanisms for these effects of the cytoplasm on MT dynamics. To distinguish between direct and indirect mechanisms, we considered two indirect mechanisms: (1) osmotic stress response pathways (such as regulation through phosphorylation), and (2) regulation by MT regulators at the MT plus end.

First, we tested whether MT stabilization is a downstream effect of an osmotic stress response pathway. Cells respond to osmotic stress by activating kinases that alter metabolism and gene expression. The MAP kinase Sty1 (Hog1, p38 ortholog) is a master integrator of multiple stress pathways (Perez and Cansado, 2011). However, in *sty1*Δ cells, sorbitol-mediated hyperosmotic shifts still caused a dampening of MT dynamics (Sup. Fig. 6) very similar to the decrease observed in wild-type cells (Fig. 1), as previously observed (Robertson and Hagan, 2008). Other triggers of the Sty1 stress pathways, such as latrunculin A, do not produce MT “freezing” (Daga et al., 2006). Thus, “freezing” of the MT network is not a downstream response of Sty1-dependent stress pathways.

We next explored the role of MT regulatory proteins at MT plus ends (+TIPs), such as MT polymerases and depolymerases. +TIPs could be affected by osmotic shifts through Sty1-independent pathways that alter their activities, affinities, phosphorylation states, etc. In fission yeast, the major classes of +TIPs are represented by the EB-family protein Mal3, which binds the GTP cap and the XMAP215- family polymerase Alp14 (XMAP215) (Akhmanova and Steinmetz, 2010; Al-Bassam et al., 2012; Busch and Brunner, 2004). In *mal3Δ* and *alp14Δ* mutant cells, hyperosmotic shifts caused a dampening of MT dynamics similar to that observed in wild-type cells (Sup. Fig.7). Therefore, the dampening is not dependent on Mal3 or Alp14. Indeed, because Mal3 is required for the recruitment of many other +TIP proteins, the *mal3Δ* mutant will cause significant disruption of the entire +TIP network—and yet MT polymerization and depolymerization still “froze” during hyperosmotic shifts. We next asked how hyperosmotic shifts impacted the localization of +TIPs. We imaged Alp14-GFP cells and observed that Alp14-GFP was maintained at the MT plus ends during hyperosmotic shifts. In contrast, Mal3-GFP localization at MT plus ends decreased (Sup. Fig. 8), consistent with the hydrolysis of the GTP cap while MTs are “frozen” (Guesdon et al., 2016), which is consistent with the prompt catastrophes we observed following the reversal of hyperosmotic shifts (Fig. 1 B and Sup. Fig. 2).

Taken together, our observations in mutant cells argue that the acute effect of increased cytoplasm concentration on MT dynamics observed here is not caused by indirect mechanisms such as the osmotic stress response or MT regulators. Rather, MT polymerization and depolymerization may be directly affected by the physical properties of the cytoplasm.

### Cytoplasmic properties modulate the motion of nanoparticles

In order to understand how cytoplasmic properties could affect a biochemical reaction like MT polymerization, we next set out to physically characterize the cytoplasm of fission yeast cells and examine how changes upon osmotic shifts affect MT dynamics. We predicted that if the cytoplasm affects MTs through physical means, then it should also affect other intracellular components not related to MTs. First, we performed microrheology experiments to assess the effects of the cytoplasm on diffusive-like motion of nanoparticles the size scale of large macromolecules. As probes, we used genetically-encoded multimeric proteins (GEMs) which assemble into spherical nanoparticles of defined sizes (Delarue et al., 2018). We expressed GEMs tagged with the fluorescent protein Sapphire in fission yeast (Methods) and imaged them at 100 fps to analyze their motion (Fig. 4 A). We analyzed GEMs of 20 and 40 nm in diameter, which are similar in size as ribosomes and of similar scale to the diameter of the MT plus end (Delarue et al., 2018). Mean squared displacement (MSD) plots revealed that the movements of the GEMs were sub-diffusive (anomalous diffusion exponent α < 1) (Sup. Fig. 9), as observed in other cell types (Delarue et al., 2018), consistent with motion being restricted by a heterogeneous meshwork of organelles and macromolecules (Luby-Phelps et al., 1986). Their diffusive-like motion at short time scales allowed us to estimate the effective diffusion coefficient (D_eff_) (see Methods and Delarue et al., 2018). GEMs D_eff_ was size-dependent: larger GEMs (40 nm, D_eff_ = 0.33 +/- 0.14) diffused more slowly than the smaller GEMs (20 nm, D_eff_ = 0.53 +/- 0.23) (Fig. 4 B), consistent with previous rheological observations in other cell types (Luby-Phelps et al., 1986; Moeendarbary et al., 2013). Thus, the cytoplasm of fission yeast appears broadly similar to cytoplasm from other eukaryotes. Interestingly, the effective diffusion coefficients of the 40-nm-GEM in fission yeast (∼ 0.3 μm^2^/sec) is similar to what was reported in budding yeast (∼ 0.3 μm^2^/sec) but slower than in mammalian cells (∼ 0.5 μm^2^/sec) (Delarue et al., 2018) perhaps revealing intrinsic differences in the level of crowding or the structure of the cytoplasm among these cell types.

**Figure 4:**
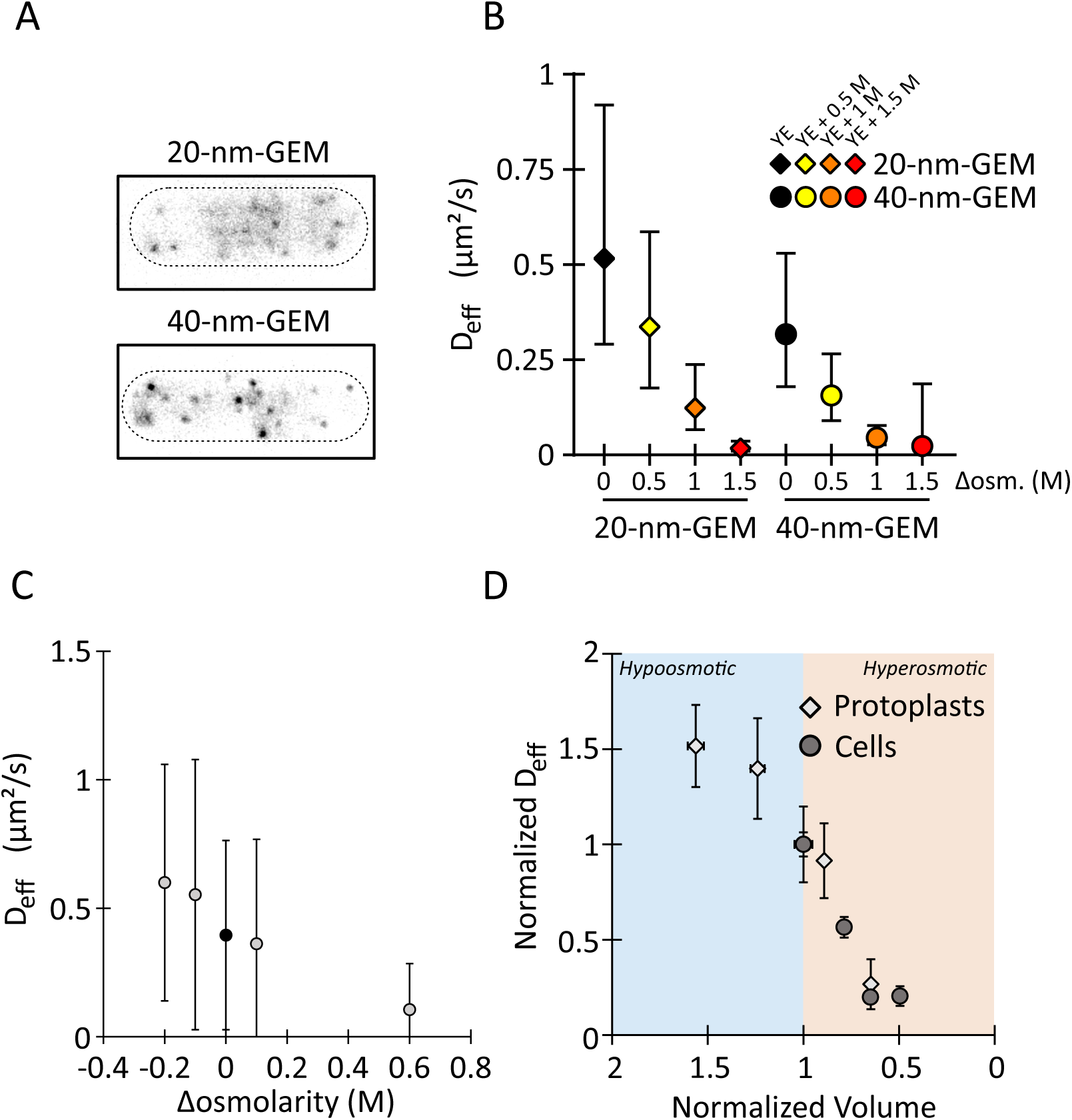
**Nanoparticle’s diffusion rate scales with cytoplasm concentration.** (A) Representative images of GEMs in fission yeast cells. Scale bar, 5 μm. (B) Hyperosmotic shifts decrease the effective diffusion coefficients of GEMs of indicated sizes in yeast cells. Values are AVG +/- standard deviation. Data come from three experiments, n > 1000 trajectories, and n > 49 cells. Concentrations reflect sorbitol concentrations in the medium. (C) Effective diffusion coefficients of 40-nm GEMs in yeast protoplasts as a function of sorbitol. AVG +/- standard deviation. Data come from at least two experiments and (left to right) n = 643 / 411 / 304 / 488 / 162 trajectories. (**H**) Diffusion rate of the 40-nm GEMs in yeast cells (circles) and yeast protoplasts (diamonds) normalized to the iso-tonic condition as a function of the normalized volume (see Methods). The rate of diffusion through the cytoplasm is faster in hypo-tonic conditions (blue shading) and slower in hyper-tonic conditions (orange shading).

Having characterized GEMs diffusion in untreated cells, we then measured the impact of osmotic shifts on the diffusion of the two different sized particles. As expected, the impact was size-dependent. At 1 M sorbitol, 40-nm GEMs were effectively immobile, while 20-nm GEMs still diffused detectably (Fig. 4 B). At 1.5 M sorbitol, GEMs of both sizes were effectively immobile (Fig. 4 B). We noted that D_eff_ decreased linearly with increasing sorbitol, particularly for the 20 nm GEM particles.

To compare effects of hypoosmotic and hyperosmotic shifts, we next analyzed GEMs in protoplasts. To establish the baseline osmolarity, we confirmed that at 0.4 M sorbitol, D_eff_ values for GEMs in protoplasts were similar to those in cells with intact cell walls (Fig. 4 C), indicating that this sorbitol concentration is the isotonic point, consistent with volume measurements. Hypoosmotic shifts increased D_eff_, while hyperosmotic shifts decreased D_eff_ (Fig. 4 C). Despite the potential complexity of the cytoplasm, these GEMs data show that the diffusion of GEMs scales inversely with cytoplasm concentration (Fig. 4C). Comparison of GEMs D_eff_ and MT rates data (Fig. 2E-F and Fig. 4C) show the same general trend that they all scale inversely with cytoplasmic concentration. These observations in yeast cells and protoplasts suggest that the physical properties of cytoplasm change with cytoplasm concentration in ways that alter the diffusion of spherical particles such as GEMs as well as MT dynamics.

### Microtubule dynamics scale with tubulin diffusion in cells

We next directly assessed the effects of osmotic shifts on the diffusion of αβ-tubulin. We measured the diffusion of soluble tubulin by fluorescence loss in photobleaching (FLIP) experiments. In FLIP, the fluorescence intensity in a whole cell is measured while a small region of the cytoplasm is repeatedly photobleached. The rate at which whole-cell fluorescence decreases over time can be used to estimate the diffusion coefficient of a fluorescent protein (Fig. 5 A) (Ishikawa-Ankerhold et al., 2012). As a probe for soluble α/β-tubulin dimers, we used Atb2-GFP (α-tubulin 2) expressed from the native chromosomal locus (Sato et al., 2009). In order to prevent MT dynamics from confounding the measurement, cells were treated with the MT inhibitor methyl benzimidazol-2-yl-carbamate (MBC) to depolymerize MTs (Fig. 5 B). To estimate the tubulin diffusion coefficient from the continuous loss of fluorescence, we designed a 1D stochastic model of tubulin diffusion that assumes a single diffusing species (Methods). Comparison of the model predictions with our experimental data (Fig. 5 B) yielded an estimated diffusion coefficient of GFP-tubulin of *D* = 7 μm^2^ s^-1^ in control cells, which is very close to the value of ∼6 μm^2^ s^-1^ obtained in pioneering experiments in sea urchin (Salmon et al., 1984) and in PTK2 cells (Wang et al., 2004). Our estimated diffusion coefficient decreased following a hyperosmotic shift, dropping to D = 4 μm^2^ s^-1^ at 1 M sorbitol and D = 1.5 μm^2^ s^-1^ at 1.5 M sorbitol (Fig. 5 C). A linear relationship emerged, namely wherein the MT polymerization and depolymerization rates correlate linearly with the tubulin diffusion coefficient (Fig. 5 D). Taken together, these data suggest that tubulin diffusion is modulated by the physical properties of cytoplasm and is likely to contribute to the observed changes in MT polymerization and depolymerization rates upon osmotic shifts.

**Figure 5:**
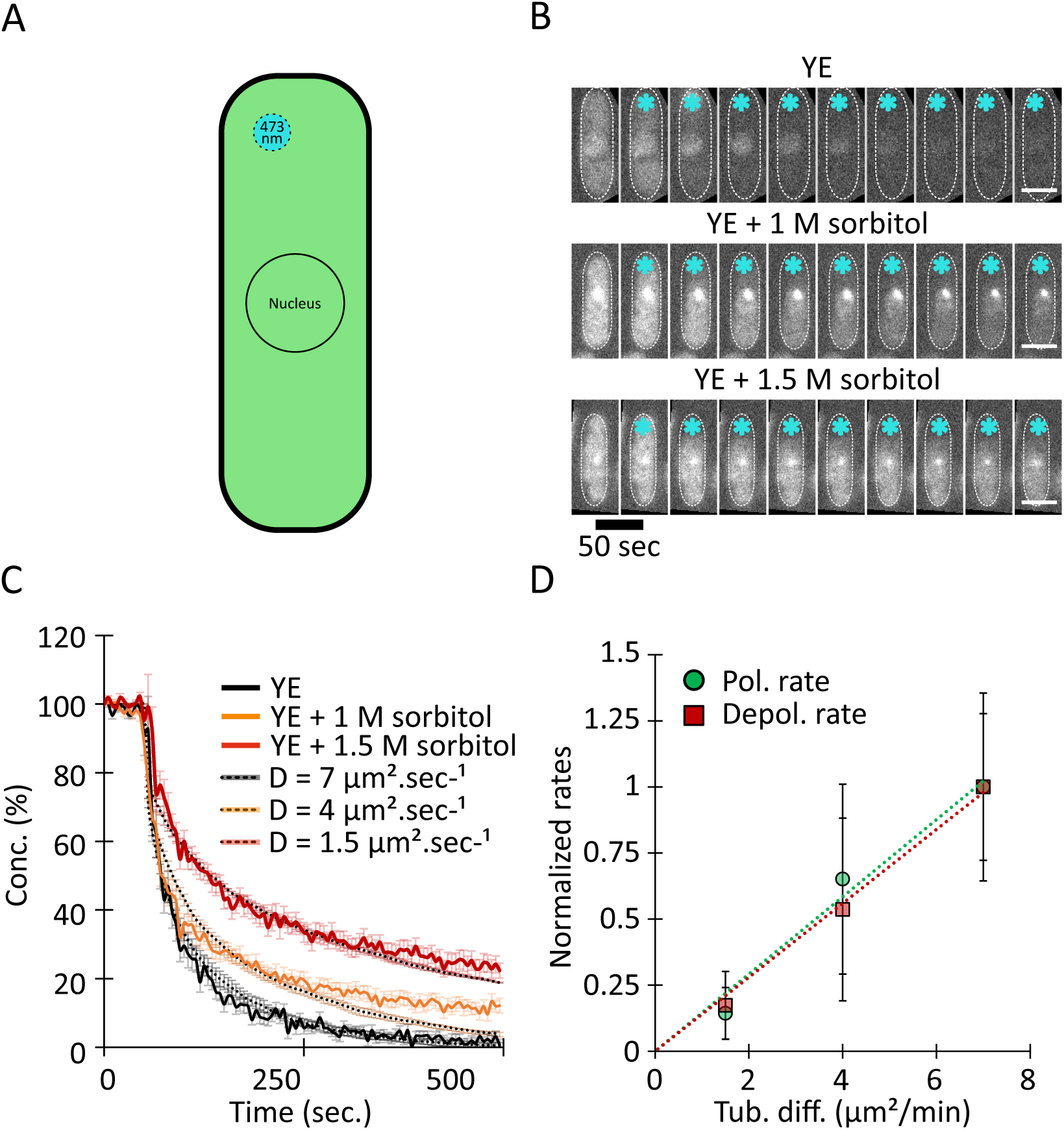
**Hyperosmotic shifts decrease the diffusion rate of soluble tubulin.** (A) To measure the diffusion of soluble GFP-tubulin, we used FLIP. Cells were exposed repeatedly to a focused laser beam (∼1 μm) near one cell tip and GFP fluorescence intensity was measured (see Methods). (B) Fluorescence decay of cells expressing GFP-tubulin in a representative FLIP experiment after hyperosmotic shock at the indicated sorbitol conditions. Interphase cells in which MTs were depolymerized with 25 μg/ml methyl benzimidazol-2-yl-carbamate (MBC) were photobleached using a focused laser (blue stars). Scale bars, 4 μm. (C) Loss of fluorescence intensity in cells from three osmotic conditions and the corresponding tubulin diffusion rate. Values (AVG +/- standard deviation) were normalized to initial intensity and expressed as concentrations (%). Data are n = 46 / 29 / 29 cells, for YE alone, YE + 1 M sorbitol, and YE + 1.5 M sorbitol, respectively, from three independent experiments. Dashed lines denote predictions from simulations of a 1D model (see Methods) for various values of diffusion; these predictions were used to estimated diffusion rates from our experimental data. Simulation values are AVG +/- standard deviation for 5 simulations. (D) Rates of MT polymerization and depolymerization in yeast cells as a function of tubulin diffusion rate. Data come from Figures 1 and 4. Correlations between diffusion and polymerization rate (P = 0.008) and depolymerization rate (P = 0.001) are significant according to Pearson’s correlation test.

### Cytoplasm viscosity increases in hyperosmotic shifts

In the ideal models of diffusion, namely the purely Brownian motion of a spherical particle in solution, the diffusion coefficient scales linearly with (1) the temperature and inversely with (2) the radius of the particle, and (3) the dynamic viscosity of the solution (Einstein, 1905). Because intracellular diffusion is complex and potentially driven by active processes, we sought to test whether viscosity of the cytoplasm changes with its concentration. We estimated fluid-phase viscosity of the fission yeast cytoplasm using time-resolved Fluorescence Anisotropy Imaging (tr-FAIM) (Siegel et al., 2003). This method measures the Brownian rotational movement of a fluorescent dye (fluorescein) in the cytoplasm of living cells, assessing viscosity at the Ångström size scale of the dye (see Methods; Fig. 6). Higher viscosity leads to lower rotation rate of the dye and a slower depolarization (Fig. 6 B). Fitting and extraction of the rate constants (Fig. 6 C) and comparison to the calibration curve (Fig. 6 D) yielded a viscosity value at 22°C of 5 +/- 1.2 cP (Fig. 6E) which is in line with the broad range of previous viscosity measurements for eukaryotic cytoplasm (range from 1 to 50 cP) (Obodovskiy, 2019). This value suggests that at 22°C the fission yeast cytoplasm has a viscosity similar to that of 43% (v/v) glycerol in water. For cells treated with 1.5M sorbitol, viscosity of the cytoplasm was 9.8 +/- 2 cP (Fig. 6 E), corresponding to 54 % (v/v) glycerol in water at 22°C. Thus, the viscosity of the cytoplasm increases with its concentration upon a hyperosmotic shift, qualitatively consistent with the effects on translational diffusion rates of GEMs and tubulin dimers.

**Figure 6:**
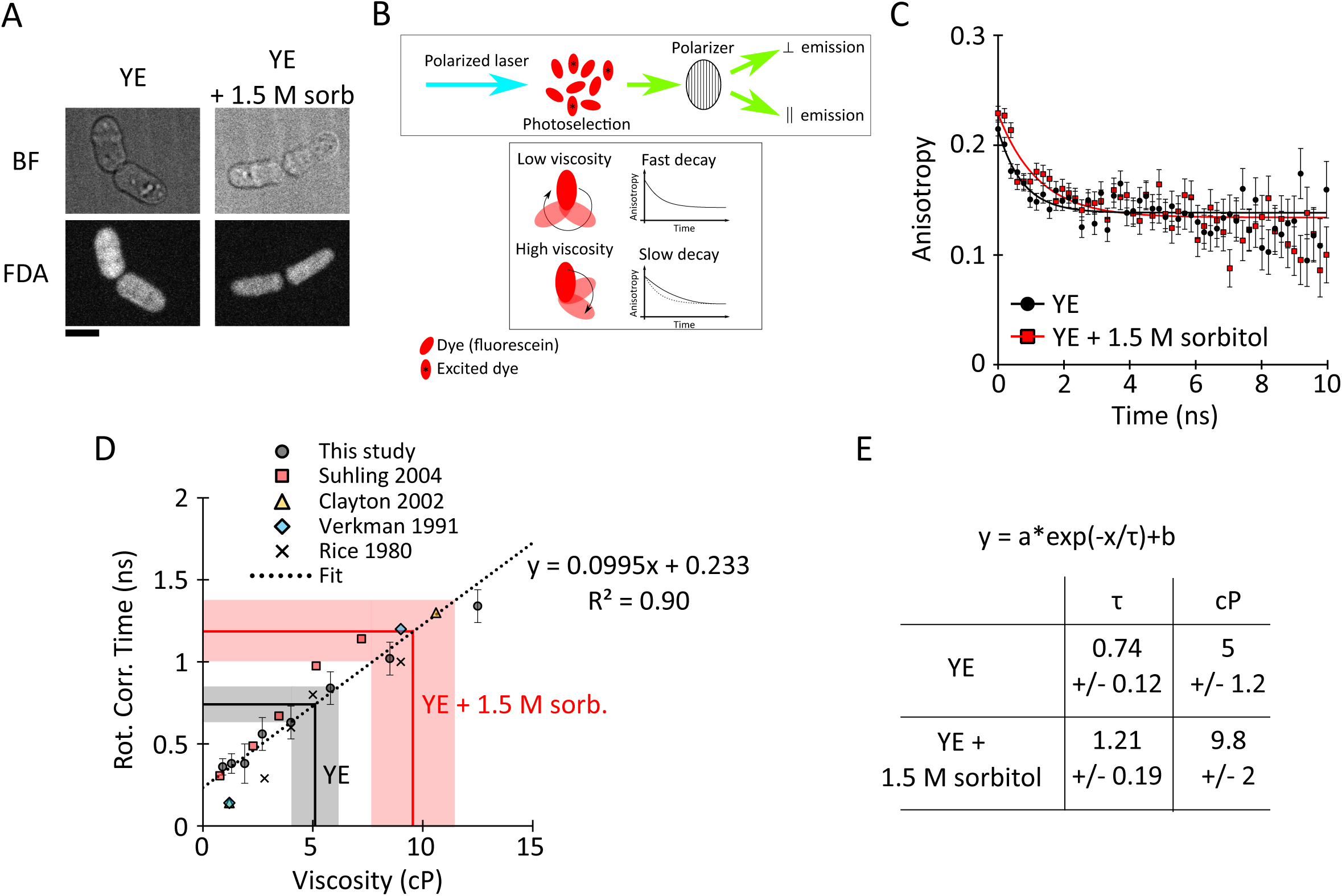
**Hyperosmotic shocks increases the viscosity of the cytoplasm in yeast.** (A) Image of cells labelled with 100 μM fluorescein diacetate (FDA) in YE and YE + 1.5 M sorbitol. Bar = 5 μm. (B) Schematic illustrating the principle of the time-resolved fluorescence anisotropy imaging (TR-FAIM) experiment. (C) Anisotropy decay of fluorescence in wildtype fission yeast cells in YE media (black circles) and YE + 1.5 M sorbitol (red squares) (AVG +/- standard deviation). The black and red plain lines represent the best fits. Differences are significant according to a F-test, p-value = 0.0003. (D) Calibration plot of fluorescein rotational correlation time in solution of different viscosity. Data from this study (black circles) and others were used. The linear regression represents the best fit. The equation was used to convert the rotational correlation time from the fits in (B) to viscosity estimates. (E) Summary of the viscosity values obtained in yeast cell in YE and YE + 1.5 M sorbitol.

### Viscosity is sufficient to explain the effects of cytoplasmic concentration

In order to isolate the effects of viscosity on MT dynamics, we reconstituted MT dynamics *in vitro* in the presence of glycerol using a well-established assay (Gell et al., 2011) (Fig. 7 A). Glycerol is a small molecule that increases viscosity in the solution without significant crowding effects. Although glycerol has been long known to stabilize MTs in bulk (Keats, 1980), its inhibitory effects on the growth rate of individual MTs *in vitro* was only recently shown (Wieczorek et al., 2013). The effect of glycerol on MT depolymerization however has not been analyzed. We took advantage of Interference Reflection Microscopy (IRM) (Mahamdeh and Howard, 2019) to image MTs at 0.5 fps for extended periods, allowing a more accurate quantification of depolymerization rates as compared to imaging fluorescently-labeled MTs. A range of glycerol concentrations was added to the reconstitutions to produce viscosities from 0.9 to 1.9 cP. MT polymerization rates decreased linearly in a dose-dependent manner with increasing viscosity, similar to what was previously shown (Wieczorek et al., 2013) (Fig. 7C). Strikingly, MT depolymerization rates also decreased with glycerol addition (Fig. 7D). Thus, the influence of cytoplasmic properties on MT polymerization and depolymerization rates that we observed *in vivo* was reproduced *in vitro* by modulating a single parameter, viscosity.

**Figure 7:**
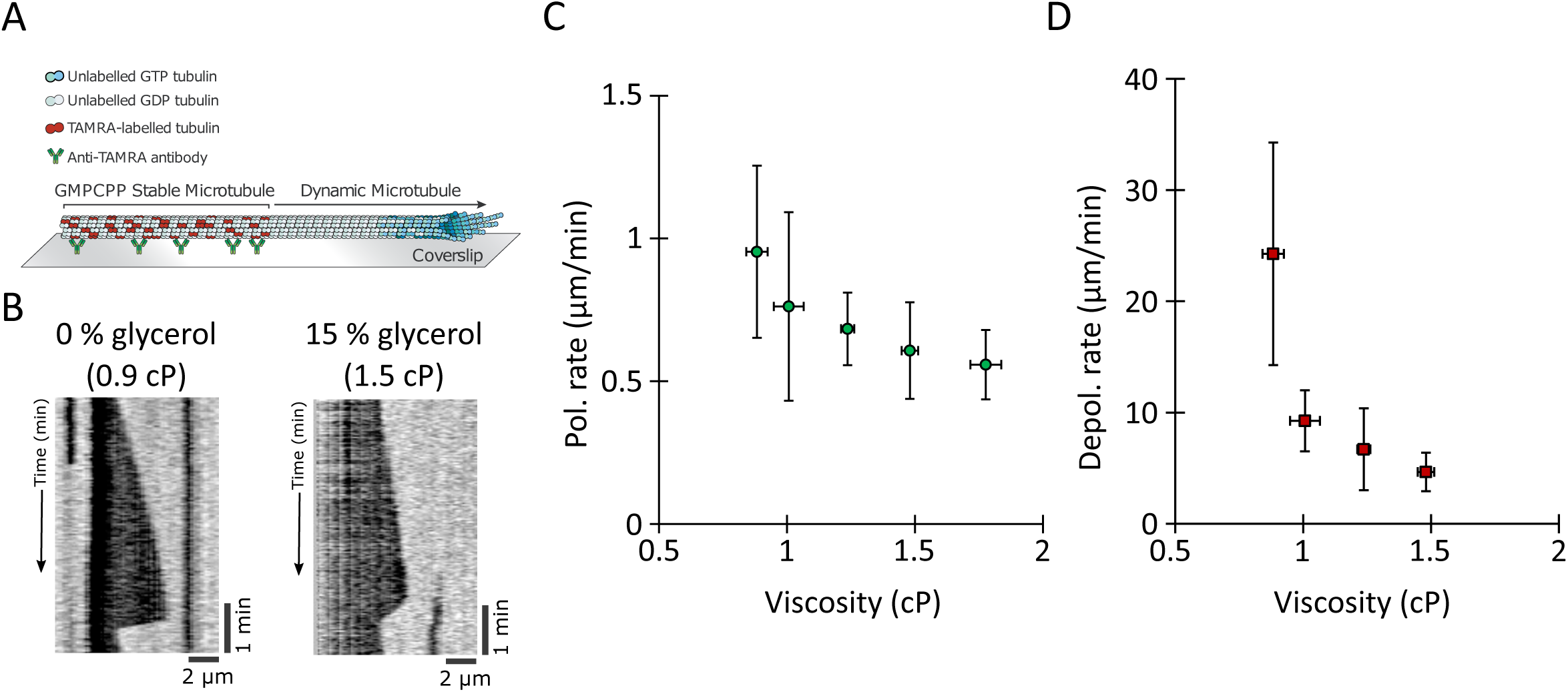
**Increasing viscosity dampens rates of MT polymerization and depolymerization.** (A) Schematic of the *in vitro* reconstituted system for MT dynamics measurement. (B) Representative kymographs of MT grown in BRB80 and BRB80 + 15 % glycerol to increase the viscosity. Polymerization and depolymerization rates are slower at higher viscosity. (C) MT polymerization rates (green circles) and (**D**) MT depolymerization rates (red squares) were measured in MTs in solutions of varying viscosity. Data represent 3 repetitions with n ≥ 70 MTs per condition.

To compare these *in vitro* results with the *in vivo* findings quantitatively, we plotted depolymerization rate as a function of polymerization rate for all the conditions we studied (Fig. 8 A). Our *in vivo* and *in vitro* experiments paint a consistent picture: rates of MT polymerization and depolymerization are linearly correlated, indicating a conserved ratio of polymerization to depolymerization (Fig. 8 A). The ratio is different in each experimental case, presumably due to the specific conditions of each case (presence of MAPs, tubulin isoform properties, tubulin concentration, temperature, etc.). Nonetheless, the ratio is maintained when MT dynamics are perturbed, either by changes in cytoplasm concentration *in vivo* or changes in viscosity *in vitro*. By normalizing the rates from each model to the value in the unperturbed condition, all of the data from *in vivo* and *in vitro* experiments strikingly collapsed onto the same slope (Fig. 8 B). This relationship shows that viscosity and the concentration of the cytoplasm affect a fundamental, conserved property of MT polymerization and depolymerization in a similar manner.

**Figure 8:**
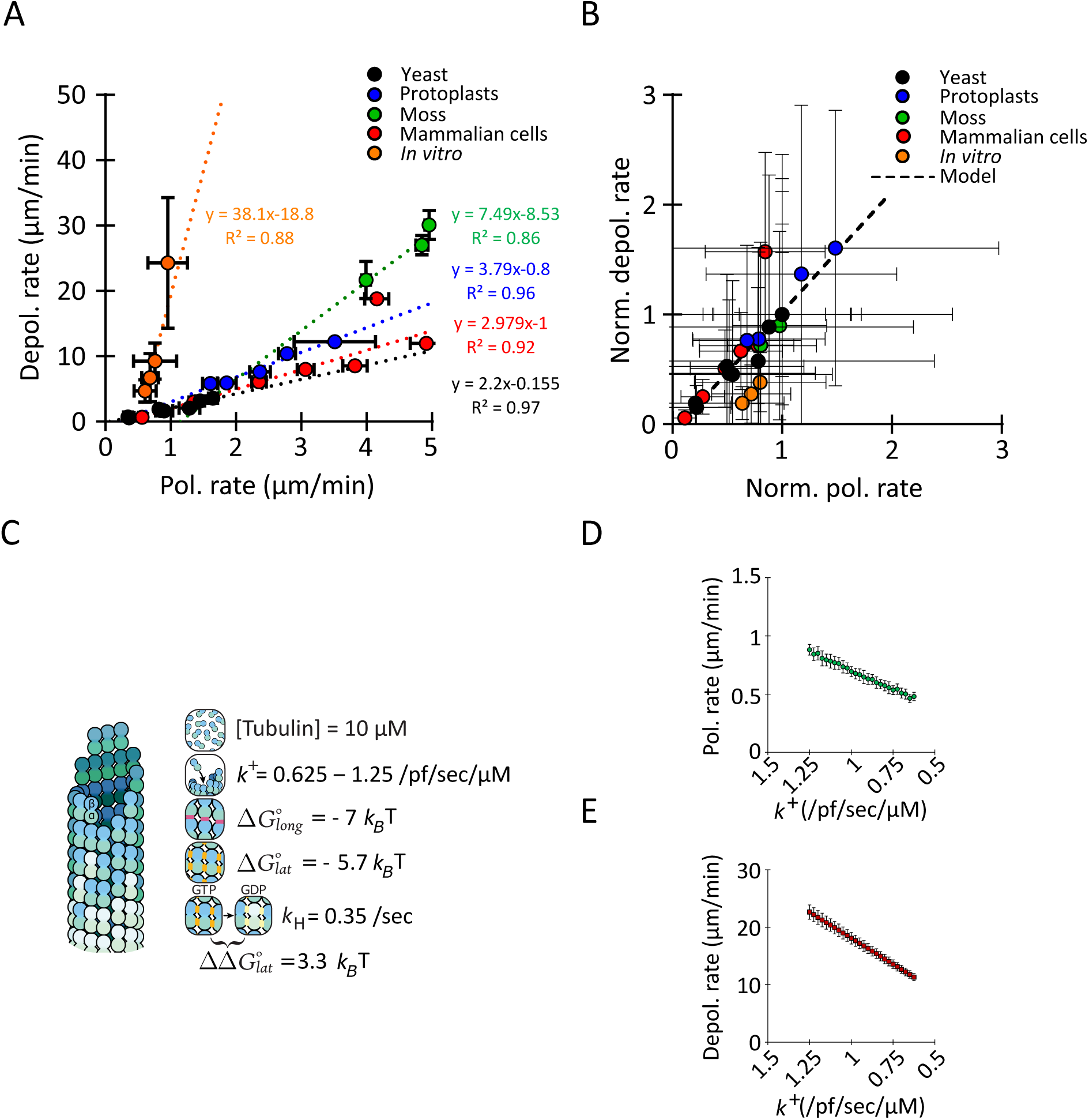
**The ratio of the rates depolymerization and polymerization is conserved when viscosity is increased.** (A) Scatter plot of the observed depolymerization rates versus the observed polymerization rates for the experiments in this study (yeast cells (black circles), yeast protoplasts (blue circles), moss (green circles), ptk2 cells (red circles), *in vitro* (orange circles)). Each model presents a certain ratio (the slope of the regression) of depolymerization rate to polymerization rate, but this ratio (the slope) is conserved when viscosity is increased *in vitro* and when cytoplasm concentration is manipulated by osmotic shocks (yeast, protoplasts, moss and Ptk2 cells) *in vivo*. AVG +/- SEM (for some data points the error bars are smaller than the point). (B) Cytoplasm and viscosity have similar effects of MT polymerization and depolarization rates. Graph shows the relationship between normalized depolymerization rates and normalized polymerization rates for all the experimental conditions and for the model (described in C). AVG +/- propagated error. All the systems (yeast, protoplast, moss, Ptk2, and *in vitro*) have the same slope than the model (see Methods). (C) Schematic of the parameters used to model MT dynamic instability. Changing the association rate constant *k*^+^ was used to model the effect of viscosity. For more details see Methods. (D) Polymerization rate (apparent on-rate) and (**E**) depolymerization rate (apparent off-rate) as a function of the association rate constant in the model. Note that the x-axis are inverted.

Using a 6 parameters model for MT dynamic instability (Fig. 8 C) (Hsu et al., 2020) that reproduces the core behaviors of polymerization, catastrophe, and depolymerization (Odde, 1997), we could reproduce these data by modulating the tubulin association rate constant *k^+^* as a proxy for the effect of viscosity. Indeed, *k^+^* describes the rate at which intermediate complexes form, and viscosity, by limiting molecular motion, should reduce it. All other parameters, notably the ΔG’s of tubulin-tubulin bonds, remained constant. The model predicts that polymerization rates decrease in a linear manner with decreasing k^+^, as expected from fewer binding events (Fig. 8 D). Notably, the model also predicts that depolymerization rates also scale linearly with *k^+^* (Fig. 8 E), because when association rate constants change, dissociation rate constants must also change if the bond energies (ΔG_long_ and ΔG_lat_) are to remain constant (Drenckhahn and Pollard, 1986). In other words, the model predicts that both rates scale linearly with the association rate constant, explaining the collapse of the normalized data on a single slope (Fig. 8 B). Thus, this initial model shows how an increase in viscosity that causes slower molecular motion can decrease both polymerization and depolymerization rates proportionally without affecting the bond energies. Thus, MT dynamic instability in the cytoplasm is modulated by a single master variable, which is viscosity.

## Discussion

Here we used MT dynamics as a model reaction for studying how the physical properties of cytoplasm influence biochemical reactions. During hyperosmotic shifts that increase cytoplasmic concentration, MTs polymerized and depolymerized more slowly and paused more frequently (Fig. 1). Conversely, in hypoosmotic conditions, which decreases cytoplasmic concentration, MT polymerization and depolymerization rates sped up as much as 50% (Fig. 2). We provide numerous lines of evidence to indicate that these effects act directly via the physical properties of the cytoplasm, as opposed to indirect mechanism via osmotic stress pathways or MT regulatory proteins (Sup. Fig. 6 and 7). The effects were rapid (Fig. 1), reversible (Sup. Fig. 2) and scaled linearly with the concentration of cytoplasm in hypo- and hyperosmotic shocks (Fig. 2), strongly indicative of a physical response. The effects on MTs scaled with effects on the diffusive motion of GEMs and tubulin dimers (Fig. 4 and 5). A significant implication is that cytoplasm properties, such as its viscosity, set the rates of MT polymerization and depolymerization at physiological conditions. Additionally, using osmotic shocks on Ptk2 cells (Fig. 3) and on moss cells (Sup. Fig. 5) we obtain similar results suggesting that the effect of the cytoplasm concentration on MT dynamics is conserved across eukaryotes. Furthermore, we can reproduce the effect of cytoplasm concentration on MTs grown *in vitro* by increasing the viscosity of the buffer (Fig. 7). Our findings further indicate that the predominant impact of cytoplasmic concentration on MTs is through its viscosity, rather than effects of macromolecular crowding or changes in tubulin concentration. Indeed, these findings provide one of the first demonstrations how the viscosity of the cytoplasm impacts an endogenous intracellular reaction.

Viscosity is likely to impact a multitude of biochemical reactions and multi-scale conformational dynamics that drive the polymerization and depolymerization of MTs (Brouhard and Rice, 2018). For MT polymerization, viscosity may inhibit diffusive arrival and positioning of a curved, GTP-tubulin dimer to the end of a protofilament (Fig. 5). Viscosity may affect the subsequent steps in which tubulin dimers straighten and form bonds between the full complement of neighboring dimers, as well as changes in proto-filament conformation during assembly of the MT lattice. For MT depolymerization, viscosity may affect the large conformational changes in protofilaments as they peel away from the MT end to form curved structures. These structural transitions, which involve significant changes in tertiary and quaternary structure (Brouhard and Rice, 2018), are influenced by solvent interactions and hence potentially by changes in viscosity. Additional studies will be needed to determine what are specific rate-limiting reactions of MT dynamics and organization responsible for these effects of viscosity.

In summary, we use MT dynamics as an example to study how the physical properties of cytoplasm affect biochemical reactions *in vivo* and discovered that viscosity plays a key role. This works highlights the impact of cytoplasmic viscosity on the rates of intracellular reactions and may generalize to diverse processes such as actin polymerization (Drenckhahn and Pollard, 1986), the assembly of multi-subunit complexes such as protein aggregation or amyloid formation (Munishkina et al., 2004), and the folding of proteins and RNA (Dupuis et al., 2018; Hagen, 2010; Pradeep and Udgaonkar, 2006). The density and other properties of the cytoplasm are known to vary during the cell cycle, in development, aging and diseases (Neurohr and Amon, 2020). Recent findings show that budding yeast may actively regulate their viscosity in response to environmental conditions such as temperature changes through regulation of metabolites such as trehalose and glycogen (Persson et al., 2020). Therefore, it will be important to consider how physiological changes in cytoplasmic properties globally affect cytoskeletal and other cellular reactions in the living cell.

## Material and methods

### Fission yeast, media, and growth conditions

Standard methods for growing and genetically manipulating *Schizosaccharomyces pombe* were used (Moreno et al., 1991). For most experiments, cells were grown overnight in Yeast Extract (YE5S) medium (here called YE) with shaking at 30 °C to exponential phase (OD_600_ between 0.2 - 0.8). See Sup. Table. for reagents and strains details.

### Strain construction

Cassettes for expressing GEMs (20 nm, AqLs-Sapphire; 40 nm, PfV-Sapphire), were cloned into the fission yeast pREP41X expression vector from pRS306 budding yeast expression vectors (Delarue et al., 2018). Briefly, GEMs expression cassettes were amplified via PCR and inserted into pREP41X via Gibson assembly at the XhoI site. The primers used for PCR are described in the supplementary table.

### Protoplast preparation

*S. pombe* cells were grown in YE5S liquid culture at 30 °C to exponential phase, harvested, and washed with SCS buffer (20 mM citrate buffer, 1 M D-sorbitol, pH 5.8), then resuspended in SCS buffer supplemented with 0.1 g/mL Lallzyme (Lallemand, Montreal, Canada) (Flor-Parra et al., 2014). Cells were digested for 10 min at 37 °C with gentle shaking in the dark. The resulting protoplasts were gently washed twice in YE5S medium with 0.2-1 M D-sorbitol, using gentle centrifugation (2 min at 0.4 rcf).

### Imaging of fission yeast

*S. pombe* cells and protoplasts were imaged in commercial microchannels (Ibidi μ-slide VI 0.4 slides; Ibidi 80606, Ibiditreat #1.5). Channels were pre-treated with 50 μl of 100 μg/ml lectin solution for 5 min at room temperature. The lectin solution was removed by pipetting and 50 μl of cell culture were introduced. After incubation for 3 to 10 minutes to allow cells to adhere to the lectin, the cells were washed with YE5S. For hyper-osmotic shocks, the medium was manually removed from the channel via pipetting and quickly replaced with hyper-osmotic media as indicated.

*Microtubule dynamics*. Yeast cells expressing GFP-tubulin (see Table) were observed with a 488 nm excitation laser at 100 ms of exposure per z-slice (1 μm spacing, 7 slices) at 0.1 Hz, with a 60x objective (CFI Plan Apochromat VC 60XC WI) on a Nikon TI-E equipped with a spinning-disk confocal head (CSU10, Yokogawa) and an EM-CCD camera (Hammamatsu C9100-13).

*Protoplast volume*. Protoplasts were resuspended YE5S medium with 0.2-1 M D-sorbitol, then imaged with a 561 nm excitation laser at 100 ms of exposure per z-slice (0.5 μm spacing), with the 60x objective, microscope, and camera used for MT dynamics.

*GEMs diffusion*. Yeast cells were imaged with a 60x TIRF objective (Nikon, MRD01691) on a Nikon TI- E equipped with a Nikon TIRF system and a SCMOS camera (Andor, Ixon Ultra 888). Protoplasts were imaged on a Nikon TI-2 equipped with a Diskovery Multi-modal imaging system from Andor and a SCMOS camera (Andor, Ixon Ultra 888) using a 60x TIRF objective (Nikon, MRD01691). Cells were imaged at 100 Hz, in TIRF, for 10 s with a 488 nm excitation laser.

*Tubulin diffusion*. Cells were imaged at 0.2 Hz, with a 60x TIRF objective (Nikon, MRD01691) on a Nikon TI-2 equipped with a Diskovery Multi-modal imaging system from Andor and a SCMOS camera (Andor, Ixon Ultra 888). Cells were imaged using spinning disk with pinhole of 100 μm. Cells were imaged at 488 nm laser excitation; the bleaching laser was a 473 nm laser controlled by a UGA-42 Firefly (Rapp OptoElectronic).

*FDA labeling*. Cells were labeled with 100 μM fluorescein diacetate (Sigma, F7378) for 30 min at room temp with agitation. Yeast cells expressing GFP-tubulin (see Table) were observed with a 488 nm excitation laser at 100 ms, with a 60x objective (CFI Plan Apochromat VC 60XC WI) on a Nikon TI-E equipped with a spinning-disk confocal head (CSU10, Yokogawa) and an EM-CCD camera (Hammamatsu C9100-13).

### Osmotic shock experiments

*Hyperosmotic shifts*. Cells were grown in rich YE5S medium, mounted into microchannels, and then treated with YE5S containing various concentrations of sorbitol while on the microscope stage at room temperature. Loss of water after the switch of medium is almost instantaneous (<30 s). To minimize adaptation responses (Tatebe et al., 2005), imaging was initiated as soon as possible (<1 min) after adding sorbitol.

*Osmotic oscillations*. Cells were introduced into a microfluidic system (Cell Asics) as described in (Knapp et al., 2019) then the media in the chamber was oscillated; YE for 5 minutes then YE with 1.5 M sorbitol for 3 minutes. During the oscillations cells were imaged for MT dynamic measurement as described above.

*Cold treatment*. Yeast cells expressing GFP-tubulin were pelleted gently and resuspended in YE containing 0, 0.5, 1, or 1.5 M sorbitol. Each culture was split into two tubes; one was incubated for 5 min at room temperature and the other was incubated on ice for 5 min. Cells were fixed by adding 16% paraformaldehyde to the medium for a final concentration of 4%. Cell were then imaged in lectin-treated Ibidi chambers.

*Osmotic shifts of protoplasts.* After cell-wall digestion, protoplasts were gently washed twice in YE5S with 0.4 M D-sorbitol using gentle centrifugation (2 min at 0.4 rcf), then placed in the Ibidi chamber for imaging. Medium was exchanged manually with hypo- or hyper-tonic medium right before imaging. YE + 0.4 M sorbitol was close to isotonic conditions, as determined by comparing volumes and GEMs dynamics to yeast cells. Thus, protoplasts resuspended in YE + 0.2, 0.25, or 0.3 M sorbitol were in hypo-tonic conditions, while protoplasts in YE + 0.5 or 1 M sorbitol were in hyper-tonic conditions.

### Measurements of MT dynamic parameters and cell volume in yeast and protoplasts

*MT dynamics*. Measurements of MT dynamic parameters were obtained using analyses of kymographs of GFP-tubulin expressing cells. Images of individual cells were cropped and multiple MT bundles per cell were selected from maximum intensity projection of the z-stack. Kymographs were made and analyzed with the KymoToolBox plugin of ImageJ (Schneider et al., 2012).

*Volume*. The effects of sorbitol on cell volume were determined from brightfield images (Atilgan et al., 2015). Protoplast volume was measured in 3D from Z-stack fluorescence images of cells expressing markers for the plasma membrane (mCherry-Psy1) using LimeSeg, a Fiji pulg-in (Schindelin et al., 2012) (Machado et al., 2019).

### FLIP experiments

Cells expressing GFP-tubulin were mounted in microchannels and treated with 25 μg/ml methyl benzimidazol-2-yl-carbamate (MBC) for >1 min to depolymerize MTs. Cells were then subjected to repeated photobleaching with a focused 473 nm laser in a 1-μm region near the cell tip using a UGA-42 Firefly system from RapOpto. Cells of similar size (∼ 12 μm long) were photobleached in order to reduce variability in the resulting data. Fluorescence decay was followed in the half of the cell submitted to the bleaching sequence to avoid the effects of diffusion around/through the nucleus. Fluorescence decay curves were normalized and aligned to the time point preceding the activation of the bleach sequence.

A calibrated 1D Brownian model of diffusion was used to simulate the fluorescence decay in a 6 μm tube (half of a cell 12 μm in length). Bleaching rate, region size, and position were matched to the experimental setup. Particles positions were updated every 0.01 s. The decay in the number of unbleached particles in the model was read out every 5 s (matching the imaging frequency) and normalized to the total number of particles. The insensitivity of the model to the total number of particles and to the time interval used was established by changing these parameters across three orders of magnitude, without significant effect on the outputs. Decay plots for various diffusion rates in the model were compared to the experimentally measured values to obtain the estimated tubulin diffusion rates.

### Microrheology with GEMs nanoparticles

GEM fusion proteins were expressed from pREP41X-based expression vectors from the thiamine-regulated nmt1* promoter (Maundrell, 1990). Transformants containing these plasmids were maintained on EMM-leu medium. The day before imaging, cells were inoculated in EMM-leu medium containing 0.05 μg/mL thiamine to allow a low level of construct expression. These conditions generally produced a few tens of GEM nanoparticles per cell. Overexpression of the GEMs commonly produced cells with single, bright, non-motile aggregates. Cells expressing GEMs were selected for sparse numbers of labeled motile nanoparticles and imaged at 100 Hz intervals. Individual cells were cropped for analysis. Nanoparticles in each cell were tracked using the MOSAIC plugin (Fiji ImageJ), and the effective diffusion rate was determined from mean squared displacement (MSD) plots as previously (Delarue et al., 2018). Briefly, tracks shorter than 10 timepoints were excluded from the MSD analysis. The following fit was used on the first 100 ms to extract the diffusion value for trajectories longer than 10 timepoints: MSD = 4Dt. The following parameters were used for the 2D Brownian dynamics tracking in MOSAIC: radius = 3, cutoff = 0, per/abs = 0.2-0.3, link = 1, and displacement = 6.

### MT dynamics in moss cells

*Growth conditions*. Moss cells were grown as described previously (Yamada et al., 2016). Caulonemal cells were mounted in microfluidic devices at room temperature as described previously (Kozgunova and Goshima, 2019).

*Osmotic shocks*. For osmotic shock, observation medium was removed manually via aspiration with a syringe, and medium with sorbitol was subsequently introduced with a syringe.

*Microtubule dynamics*. Cells were imaged on a Nikon TI-E TIRF system with a 60x TIRF objective (Nikon, MRD01691) and a SCMOS camera (Andor, Ixon Ultra 888). MT dynamic parameters were obtained using kymographs. Kymographs were made and analyzed with the KymoToolBox plugin of ImageJ.

### MT dynamics in mammalian cells

*Culture condition*. PTK2 cells stably expressing GFP-tubulin were grown in DMEM/F12 medium supplemented with 10% fetal bovine serum (FBS) and 1% antibiotic-antimycotic (Thermofisher #15240062) (10% DMEM/F12) with 5% CO2. The day before experiments, cells were plated in 35 mm glass bottom dish (Ibidi #81218), incubated in 2 ml of medium and left overnight at 37 °C and 5% CO2 to allow them to spread properly.

*Osmotic shocks*. The media contained in the 35 mm glass bottom dish was manually replaced by 2 ml of 10% DMEM/F12 prewarmed to 37 °C containing various concentrations of sorbitol while on the microscope stage controlled with the Chamlide TC incubator (kept at 37 °C and 5% CO2). Loss of water after the switch of medium is almost instantaneous (<30 s). To minimize adaptation responses, imaging was initiated 3 minutes before the switch of medium and continued for 4 minutes after adding sorbitol.

*Microtubule dynamics*. PTK2 cells stably expressing GFP-tubulin plated in 35 mm glass bottom dish were observed with a 488 nm excitation laser at 200 ms of exposure at 0.2 Hz with a 60x objective (Plan Apo VC 60x/1.40 oil) in a Nikon TI-E equipped with a spinning disk confocal head (CSU-X1, Yokogawa) and a CCD camera (QImaging Retiga R3). The cell culture conditions on the microscope stage were controlled with the Chamlide TC incubator (kept at 37 °C and 5% CO2).

*Analysis*. Measurements of MT dynamic parameters were obtained using kymographs of PTK2 cells stably expressing GFP-tubulin. Region of interest (ROIs) of individual cells were cropped and multiple MTs per ROI were selected. Kymographs were made and analyzed with the KymographBuilder plugin of ImageJ.

### MT dynamics *in vitro*

*Tubulin preparation.* Tubulin was purified from juvenile bovine brains via cycles of polymerization and depolymerization, as described previously (Ashford and Hyman, 2006). GMPCPP-stabilized MT seeds were prepared by polymerizing a 1:4 molar ratio of tetramethylrhodamine (TAMRA, ThermoFisher Scientific) labeled:unlabeled tubulin (Hyman, 1991) in the presence of GMPCPP (Jena Biosciences) in two cycles, as described previously (Gell et al., 2011).

*MT reconstitution assay.* Dynamic MTs were imaged in a reconstitution assay from surface-bound, stabilized MT seeds (Gell et al., 2011). Cover glass was cleaned, as previously described (Helenius et al., 2006). Two silanized cover glasses (22 x 22 mm and 18 x 18 mm) were separated by multiple strips of double-sided tape on custom-machined mounts to create channels for solution exchange. Channels were prepared for experiments by flowing in anti-TAMRA antibodies (ThermoFisher Scientific) and blocking with 1% Pluronic F-127 for 20 min. Channels were rinsed three times with BRB80 before flowing in seeds and placing the chamber on the microscope stage, where the objective was heated to 32 °C with a CU-501 Chamlide lens warmer (Live Cell Instrument).

Dynamic MTs were grown from GMPCPP seeds by filling the channel with 10 µM tubulin in reaction buffer: BRB80 (80 mM PIPES-KOH [pH 6.9], 1 mM EGTA, 1 mM MgCl_2_) plus 1 mM GTP, 0.1 mg/mL bovine serum albumin, 10 mM dithiothreitol, 250 nM glucose oxidase, 64 nM catalase, and 40 mM D-glucose. Reaction buffer was prepared on ice before being flowed into the channel with a piece of filter paper. A 60% (v/v) glycerol stock solution in BRB80 was added to the indicated final concentrations. For consistency, a large aliquot of tubulin was thawed on the day of each experiment, sub-aliquoted, and stored in liquid nitrogen. A separate sub-aliquot was thawed for each individual experiment. Glycerol concentrations from 0-15% (v/v) were tested. We note that the effects of glycerol concentrations ≥20% were not measurable as no depolymerization events were observed because of the high levels of spontaneous nucleation and inhibition of catastrophes at those glycerol concentrations.

*Interference reflection microscopy*. Dynamic label-free MTs were imaged with interference reflection microscopy (IRM) as described previously (Mahamdeh and Howard, 2019).

*MT dynamics*. MT dynamics was analyzed using kymographs as described above.

*Viscosity measurements in vitro*. Viscosities of glycerol solutions were measured using an mVROC Viscometer (Rheosense) at room temperature (23°C as measured by the viscometer). Measurements were made at a flow rate of 1000 µl/min for 5 sec for three replicates of each solution. Using the expected values of viscosity for similar water-glycerol mixtures (http://www.met.reading.ac.uk/#sws04cdw/viscosity_calc.html) at 23°C and 32°C, we extrapolated the viscosity of the buffers at 32°C from the measurements at 23°C.

### Cytoplasmic viscosity measurements

*FDA labeling*. Wild-type fission yeast cells were grown in YE overnight at room temperature with agitation to below saturation. Cells were labeled with 100 μM fluorescein diacetate (Sigma, F7378) for 30 min at room temperature with agitation.

*Time-resolved fluorescence anisotropy.* Cells were imaged in a Ibidi μ-slide VI 0.4 (Ibidi 80606, Ibiditreat #1.5) treated with lectin on an inverted TCS-SP2 microscope (Leica Microsystems, Germany) while the time- and polarization-resolved intensity decay was measured on SPC-150 TCSPC boards (Becker & Hickl, Germany). The emission was split using a polarizing beam splitter. Fluorescein diacetate (FDA) was excited using a picosecond-pulsed (90 ps optical pulse width, 20 MHz repetition rate) 467 nm diode laser (Hamamatsu, Japan) through a 63X 1.2 NA water objective, with a 485 nm dichroic mirror and a 500 nm long pass filter. All measurements were performed at room temperature (22°C).

Parallel (I_∥_) and perpendicular (I_⊥_) intensity decays were used to calculate the anisotropy decay, according to

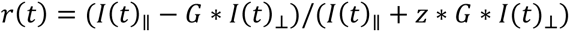

where z = 1 to account for the depolarization of the high magnitude objective and G was 0.96. Z was determined by comparing the calculated total intensity decay

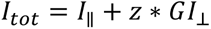

of fluorescein measured on the 63x objective and measured on a 5x objective, altering the value of z between 2 and 1 until the lifetime was an exact match (Suhling et al., 2014). G is a correction factor to account for the sensitivity of the two detectors and is calculated by taking the tail value of

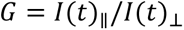

after tail-matching of the parallel and perpendicular decays by adjustment of the polarizing beam splitter.

Parallel and perpendicular intensity decays were summed over six measurements (3 measurements x 2 biological repeats) to obtain high photon counts, after ensuring negligible variation between measurements and repeats. The resulting time-resolved anisotropy decay was fit to a monoexponentially decaying model according to

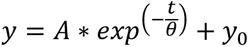

where θ is the rotational correlation time, A is the amplitude of the decay (in terms of the anisotropy decay, this value is equal to r_0_-r_ꚙ_), and y_0_ is the level to which the anisotropy decays (properly known as r). The rotational correlation time was converted into a viscosity value in cP using a calibration plot in the relevant viscosity region.

*Fluorescein calibration curve.* Methanol solutions containing various amounts of glycerol (0-40%) were prepared. Solution viscosities at room temperature (22 °C) were established previously using a rheometric expansion system rheometer (ARES) (Kuimova et al., 2008) (Steinmark et al., 2020). Fluorescein (10 μM) was added to the solutions and fluorescence was measured at room temperature to extract the rotational correlation times as described above. Note that the calibration was carried out on a 5x air objective; thus, z was taken as 2. Data from the literature (Clayton et al., 2002; Rice and Kenney-Wallace, 1980; Suhling et al., 2004; Verkman et al., 1991) were include in the calibration.

*Anisotropy decay*. Anisotropy data were fitted using GraphPad Prism. Fits used 1/STD^2^ weighting and were limited to the first 10 ns of recording. Best-fit values for each parameter and condition were obtained with a F-test. Best fits were different with p-value = 0.0003.

### Normalizations

*MT dynamics versus cellular volume*. MT dynamics and cell volume in yeast were normalized to the YE condition (no sorbitol). In protoplasts MT dynamics and volume were normalized to the 0.4 M sorbitol condition.

*GEMs diffusion versus volume*. Diffusion values and volumes were normalized to the YE (no sorbitol condition) for yeast and to the 0.4 M sorbitol condition for protoplasts.

*Polymerization rate versus depolymerization rate*. For yeast, data rates were normalized to the YE (no sorbitol) condition. For protoplasts, data were normalized to the 0.4 M sorbitol condition. For moss, data were normalized to the BCD (no sorbitol) condition. For *in vitro*, data were normalized to the BRB 80 (no glycerol) condition. For the model, data were normalized to the k_+_= 1.25 /pf/μM condition. A linear fit on the normalized data from the model gives a slope of 1.1 +/- 0.2. Normalized data from all the other systems (yeast cells, yeast protoplasts, moss cells, in vitro) collapse on the same slope according to a F-test. P values were 0.2, 0.9, 0.8, and 0.38 for yeast cells, yeast protoplasts, moss cells and *in vitro* respectively.

## Acknowledgments

We thank members of the Chang lab, and Sophie Dumont and her lab for fruitful discussions. We are grateful to Dr. Rikki Garner for countless discussions and her help setting up the stochastic model of diffusion, and to Dr. Fabrice Cordelières for the KymoToolBox ImageJ plug-in. We thank Sherman Foo and the Oliferenko lab for reagents and technical support. We are grateful to Steve Ross and Nikon for microscope support at MBL, and to the MBL communities from the Physical Biology of the Cell Course, the Physiology Course and the Whitman Investigator Program for support and advice.

## Author Contributions

Conceptualization: A.T.M. and F.C.; Methodology: A.T.M., J.L, M.G., C.H.E., C-T.H., I.E.S., K.S., G.G., L.H., G.J.B., M.T. and F.C.; Formal analysis: A.T.M., J.L, M.G., C. H. E., I.E.S; Investigations: A.T.M., C.H.E., I.E.S; Writing: A.T.M., G.J.B., and F.C.; Review editing: A.T.M., C.H.E., G.J.B., and F.C.; Visualization: A.T.M.; Supervision: A.T.M.; Project administration and funding acquisition: F.C.

## Supplemental Information

Refers to the webpage for description and access to the supplemental material.

