## Supplemental_Material for "Physical properties of the cytoplasm modulate the rates of microtubule polymerization and depolymerization"

### Supplemental Material Legends

**Supplemental figure 1: Cytoplasm concentration increases after hyperosmotic shock.**

(**A**) Normalized yeast cell volume in YE alone and in YE with increasing the indicated amounts of sorbitol. AVG ± propagated error. Data come from at least 2 experiments and (left to right) n = 269 / 90 / 76 / 38 / 64 / 42 / 24 / 51 cells, respectively.

(**B**, **C**) Averaged normalized fluorescence per cell in YE, YE + 1 M sorbitol, and YE + 1.5 M sorbitol. Yeast cells expressed (**B**) GFP-atb2 (tubulin) or (**C**) rps802-GFP (ribosome). Cellular contents increase after osmotic shocks.

(**D**) No cellular content is lost through osmotic shocks, as shown by the conservation of the total amount of fluorescence per cell estimated by multiplying the normalized volume from (**A**) by the normalized fluorescence from (**B**) and (**C**). AVG ± propagated error.

**Supplemental figure 2: The effect of sorbitol on MT dynamics is highly reversible.**

MT dynamics stops rapidly at every transition from YE to YE + 1.5 M sorbitol and restarts equally rapidly at the reverse transition. Left: (Top) Brightfield and (Bottom) fluorescence imaging of the first time point of the time-lapse. Right: Kymographs of MT dynamics in cells labeled (a), (b), and (c) at left. Cells were imaged every 5 s through 7 z-slices (1 µm spacing) for ~30 min. Images are maximum z-projections. MT dynamics are projected along the entire cell length and width. Using microfluidic CellASICS (Millipore) plates (Y04C), cells were exposed to YE + 1.5 M sorbitol 3 times for 3 min with a wash in YE medium alone for 5 min between exposures.

**Supplemental figure 3: Osmotic perturbation via sorbitol renders MTs less dynamic.**

Sorbitol exposure narrows the distributions of the rate of MT length change, indicating reductions in both polymerization and depolymerization rates. The frequency of pausing also increases upon sorbitol exposure. Positive rates correspond to polymerization events, negative rates to depolymerization events, and null rates to pauses. Left to right, n = 668 / 494 / 252 / 261 / 314 / 118 / 45 / 96 events from top left to bottom right.

**Supplemental figure 4: Hyperosmotic shock protects MTs against depolymerization by cold treatment.**

Left: Cells expressing GFP-tubulin were diluted in medium containing 0, 1, or 1.5 M of sorbitol, then incubated for 5 min at room temperature or on ice and then fixed with 3.2% paraformaldehyde (Methods). After fixation, z-stacks (7 slices, 1 µm spacing) were acquired via spinning disk microscopy (Methods). Maximum projections of representative fields of view are shown for each condition. Scale bars, 5 µm. Cold treatment depolymerized MTs in all cells in YE without sorbitol, in most cells in YE + 1 M sorbitol, and in none of the cells in YE + 1.5 M.

**Supplemental figure 5: Hyperosmotic shocks reduce MT dynamics in moss cells.**

(**A**) Representative brightfield (BF) and total internal reflection fluorescence microscopy of moss cells expressing EB1-citrine and mCh-tubulin in BCD medium without sorbitol and BCD + 0.5 M sorbitol. The bottom three rows are representative kymographs of MT polymerization events in BCD and BCD + 0.5 M sorbitol.

(**B**) MT polymerization (growth; green circles) and (**C**) depolymerization (shrinkage; red squares) rates in moss cells treated acutely with the indicated sorbitol concentrations. AVG $\pm$ standard deviation. Data come from three experiments, with (left to right) n = 216 / 157 / 112 / 27 polymerization events and (left to right) 71 / 38 / 27 depolymerization events from 3 experiments.

**Supplemental figure 6: The effect of hyperosmotic shocks on MT dynamics is independent of the Sty1 stress pathway.**

MT (**A**) polymerization and (**B**) depolymerization rates in wildtype (WT) and *sty1*Δ mutant strains in YE medium and after hyperosmotic shocks in YE medium with the indicated concentrations of sorbitol. Rates are normalized to the rates in YE. AVG $\pm$ standard deviation; $\geq$2 experiments.

**Supplemental figure 7: MT dynamics in strains lacking Mal3 and Alp14.**

Left: Representative (**A**) *mal3Δ* or (**B**) *alp14Δ* cells expressing a GFP-atb2 chimera in YE medium with the indicated sorbitol concentration. Cells were imaged with spinning disk confocal microscopy. Yellow arrows highlight dynamic MT tips and red arrows highlight "frozen" MT tips. Right: Kymographs of the cells pictured at left reveal that in YE + 1.5 M sorbitol, MT dynamics is stopped in the mutants.

**Supplemental figure 8: +TIP regulatory protein localization after hyperosmotic shock.**

Field of view of cells expressing (**A**) alp14-GFP or (**B**) mal3-GFP with atb2-mCherry in YE medium with the indicated sorbitol concentrations. Cells were imaged with spinning disk confocal microscopy. Alp14 (XMAP215) is localized at the MT plus-end in all sorbitol conditions. Mal3 (EB1) is localized at the plus-end in YE without sorbitol. In sorbitol-treated cells, Mal3 localization is less evident and seems to decrease with the amount of sorbitol.

**Supplemental figure 9: Mean squared displacement (MSD) of GEMs nanoparticles.**

Aggregated MSD of the motion of **(A)** 20-nm GEMs and **(B)** 40-nm GEMs in the cytoplasm of cells in medium with the indicated amounts of sorbitol. For each plot, the dotted and dashed lines correspond to MSD = $4Dt^{\alpha}$, where D = 0.5 $\mu m^2 s^{-1}$ and $\alpha$ = 1 (dotted line) and $\alpha$ = 0.7 (dashed line). $Log_{10}$ transformations of data in **(A-C)** appear in **(D-F)**, respectively.

**Supplemental table 1: List of strains and reagents.**

Sup. Fig. 1

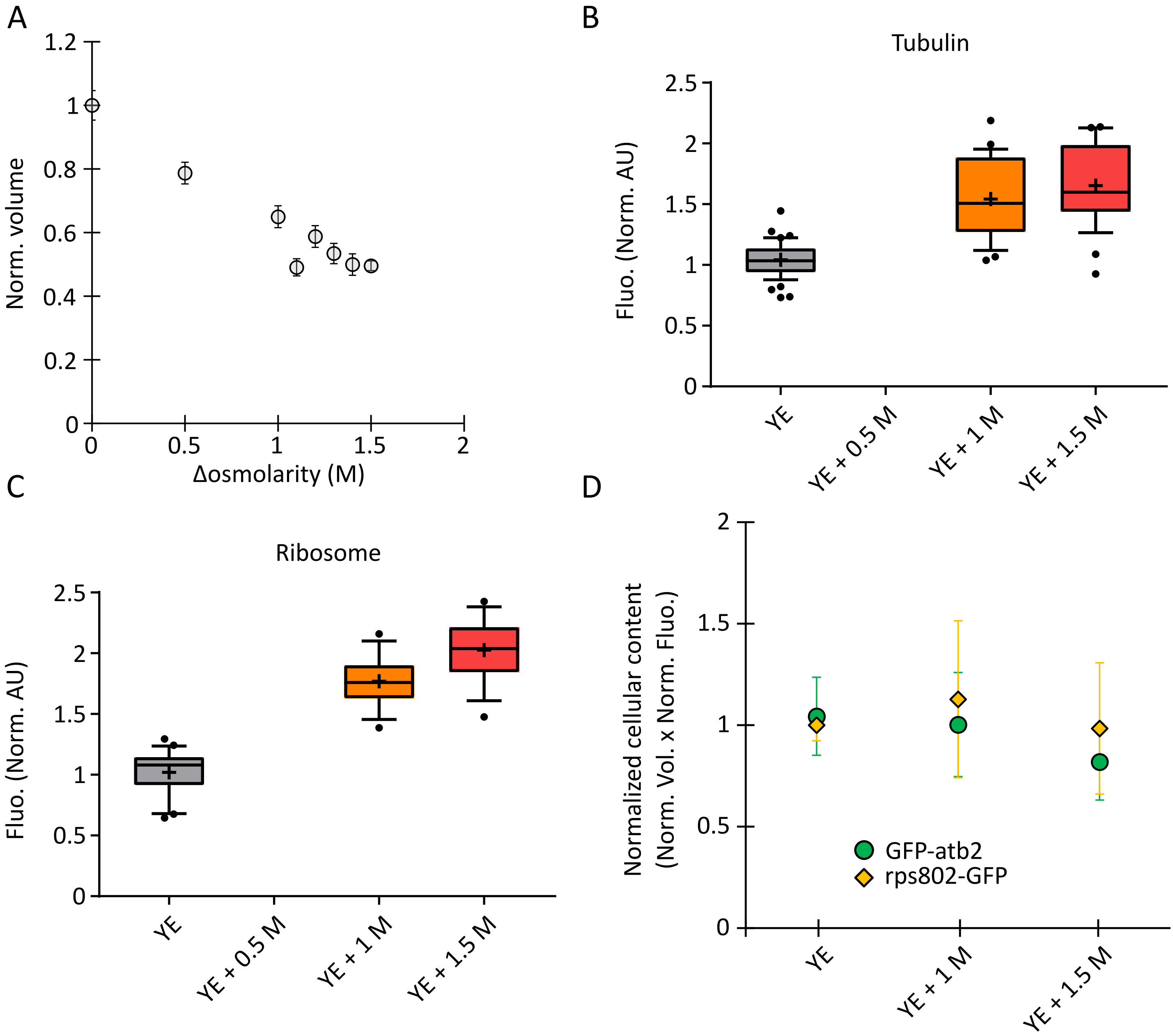
A
1.2
1
0.8
0.6
0.4
0.2
0
Norm. volume
0
0.5
1
1.5
2
Δosmolarity (M)
B
Tubulin
2.5
2
1.5
1
0.5
0
Fluo. (Norm. AU)
YE
YE + 0.5 M
YE + 1 M
YE + 1.5 M
C
Ribosome
2.5
2
1.5
1
0.5
0
Fluo. (Norm. AU)
YE
YE + 0.5 M
YE + 1 M
YE + 1.5 M
D
2
1.5
1
0.5
0
Normalized cellular content
(Norm. Vol. x Norm. Fluo.)
GFP-atb2
rps802-GFP
YE
YE + 1 M
YE + 1.5 M

### Sup. Fig. 2

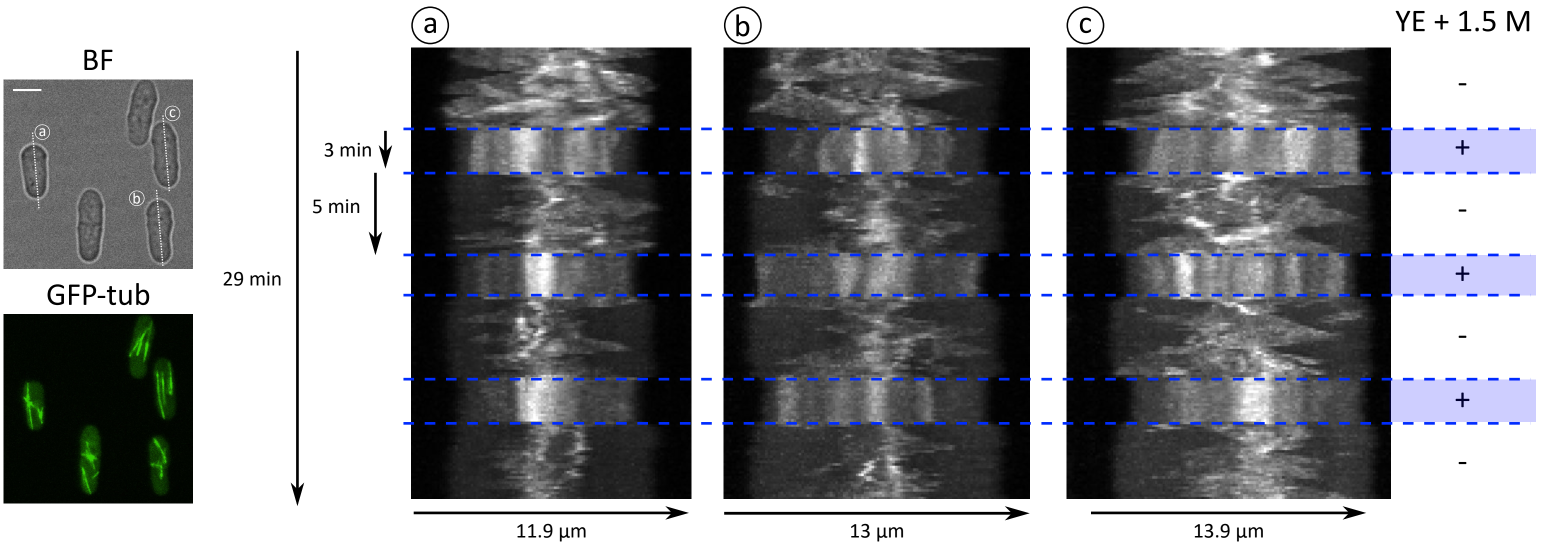

### Sup. Fig. 3

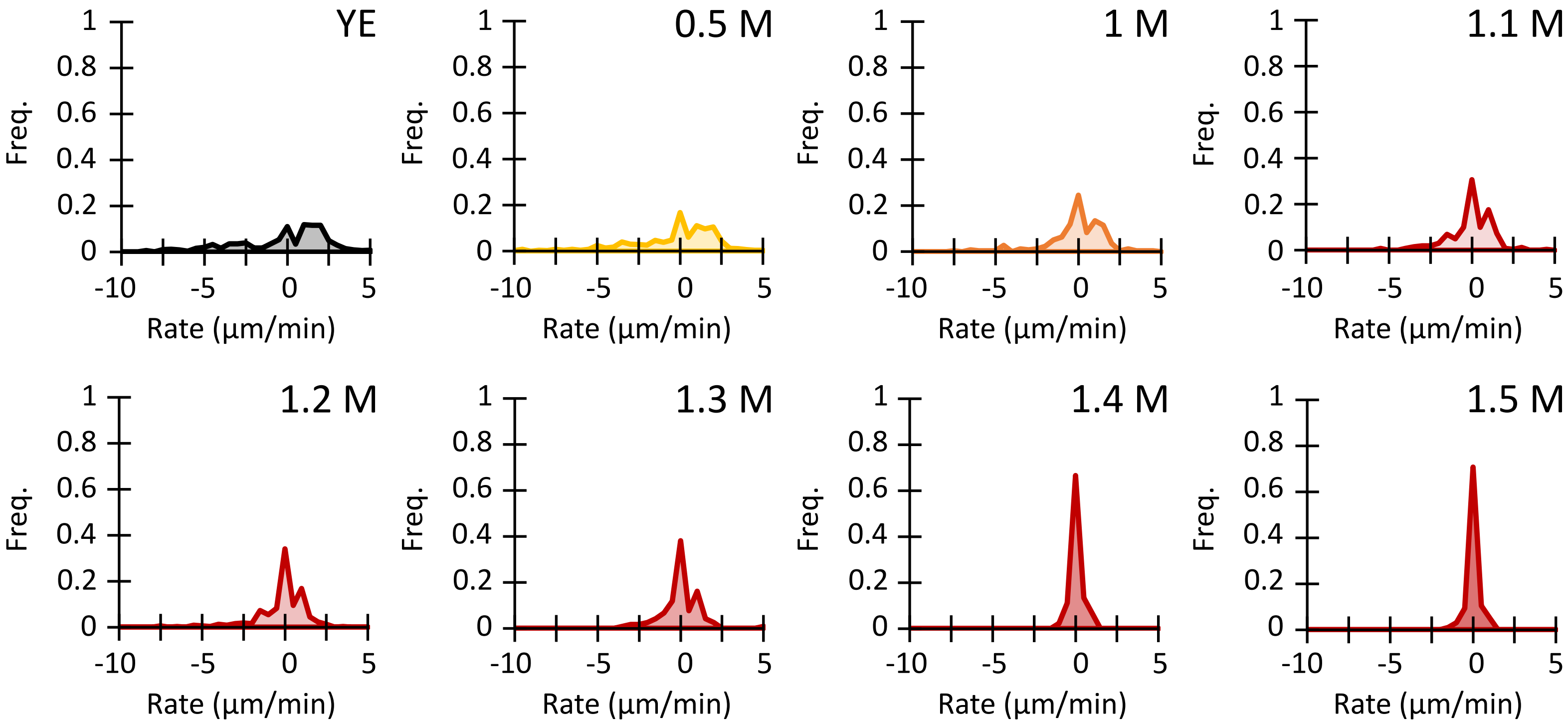

YE
0.5 M
1 M
1.1 M
1.2 M
1.3 M
1.4 M
1.5 M
Freq.
1
0.8
0.6
0.4
0.2
0
-10
-5
0
5
Rate (µm/min)

Sup. Fig. 4

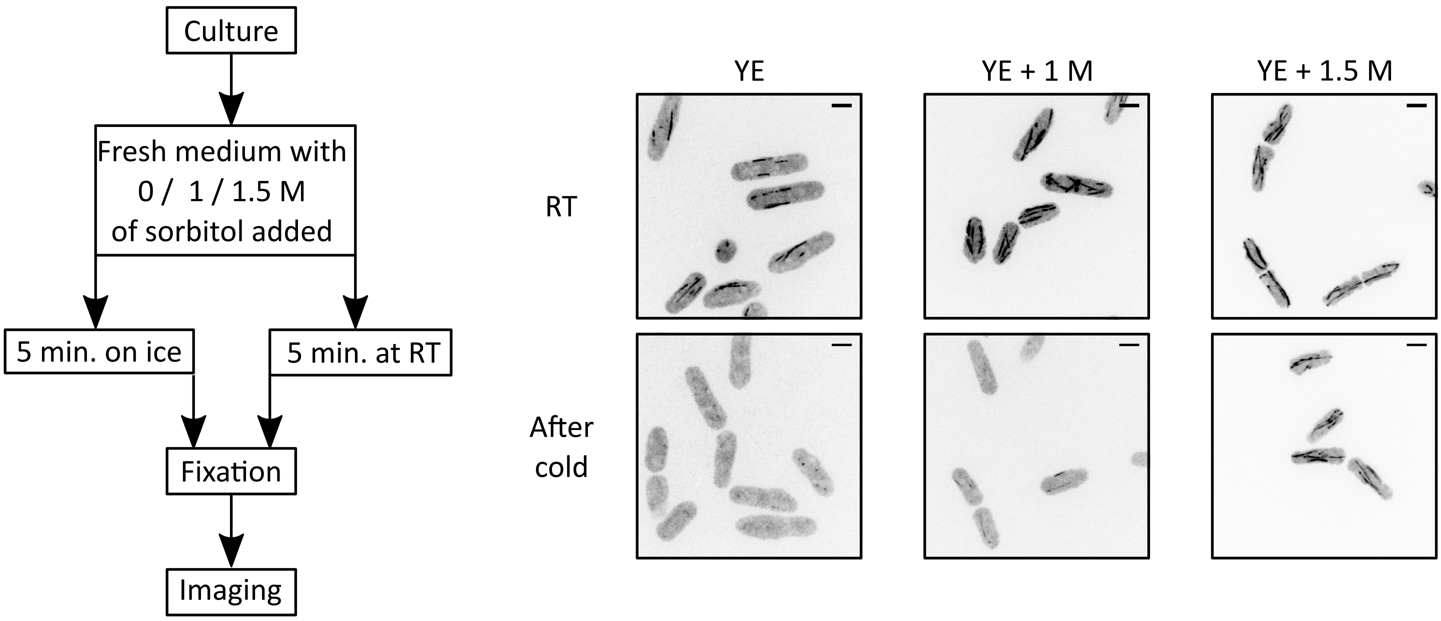
Culture
Fresh medium with 0 / 1 / 1.5 M of sorbitol added
5 min. on ice
5 min. at RT
Fixation
Imaging
YE
YE + 1 M
YE + 1.5 M
RT
After cold

### Sup. Fig. 5

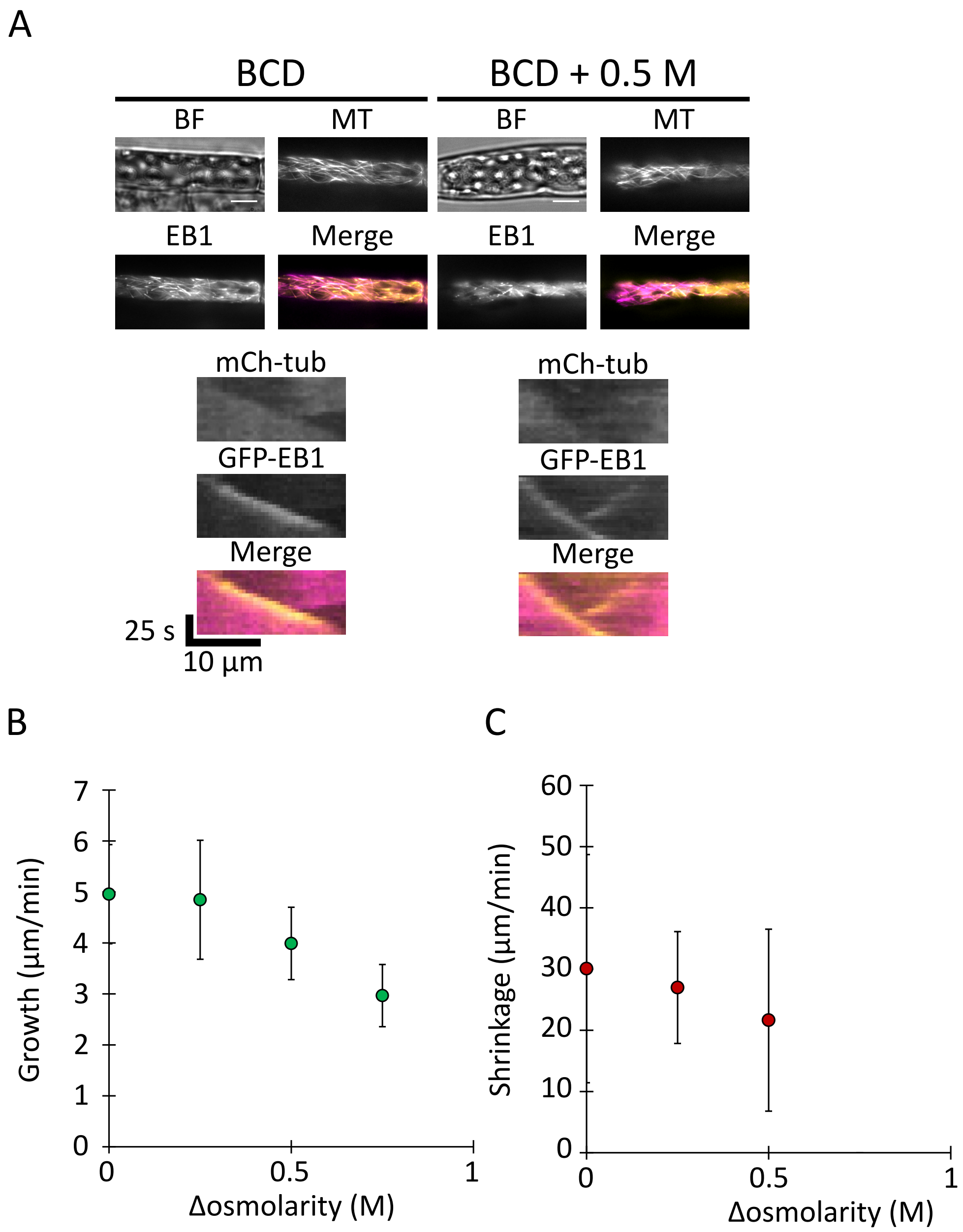
A
BCD
BCD + 0.5 M
BF
MT
EB1
Merge
mCh-tub
GFP-EB1
25 s
10 µm
B
Growth (µm/min)
0
1
2
3
4
5
6
7
0
0.5
1
Δosmolarity (M)
C
Shrinkage (µm/min)
0
10
20
30
40
50
60
0
0.5
1
Δosmolarity (M)

Sup. Fig. 6

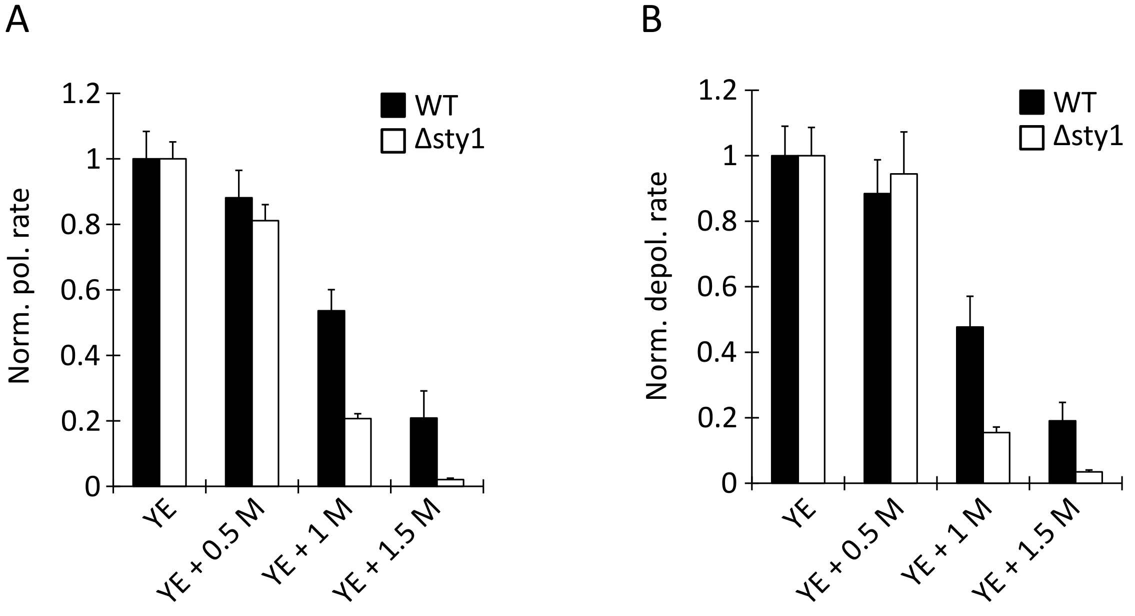
A
1.2
1
0.8
0.6
0.4
0.2
0
Norm. pol. rate
WT
Δsty1
YE
YE + 0.5 M
YE + 1 M
YE + 1.5 M
B
1.2
1
0.8
0.6
0.4
0.2
0
Norm. depol. rate
WT
Δsty1
YE
YE + 0.5 M
YE + 1 M
YE + 1.5 M

Sup. Fig. 7

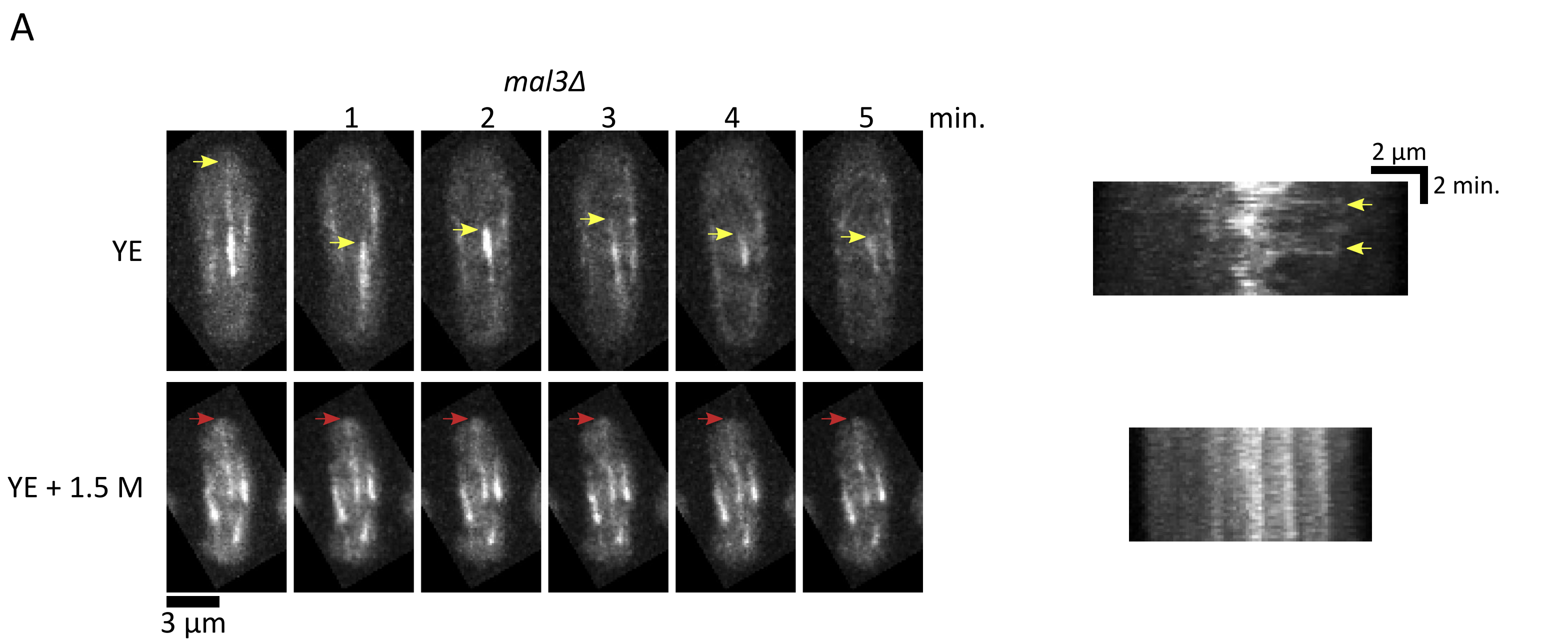
A
*mal3Δ*
1
2
3
4
5
min.
YE
YE + 1.5 M
3 µm
2 µm
2 min.

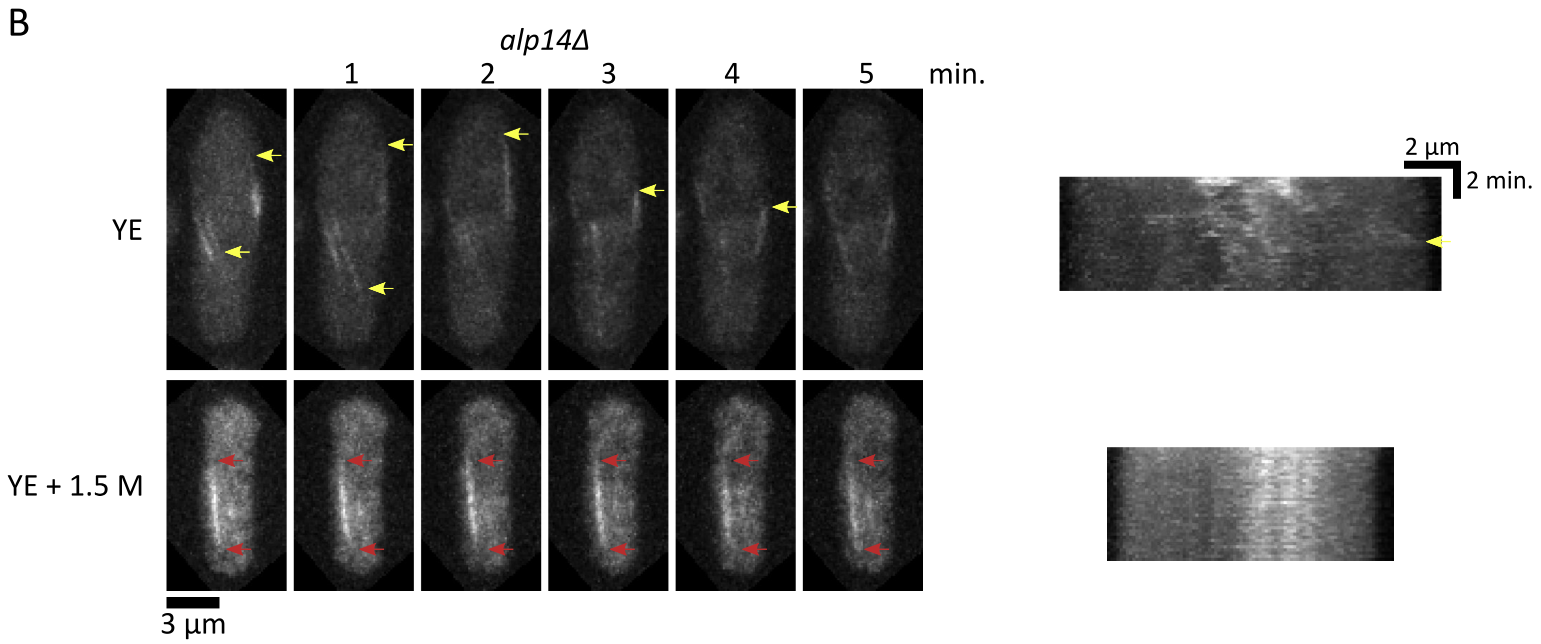
B
*alp14Δ*
1
2
3
4
5
min.
YE
YE + 1.5 M
3 µm
2 µm
2 min.

Sup. Fig. 8

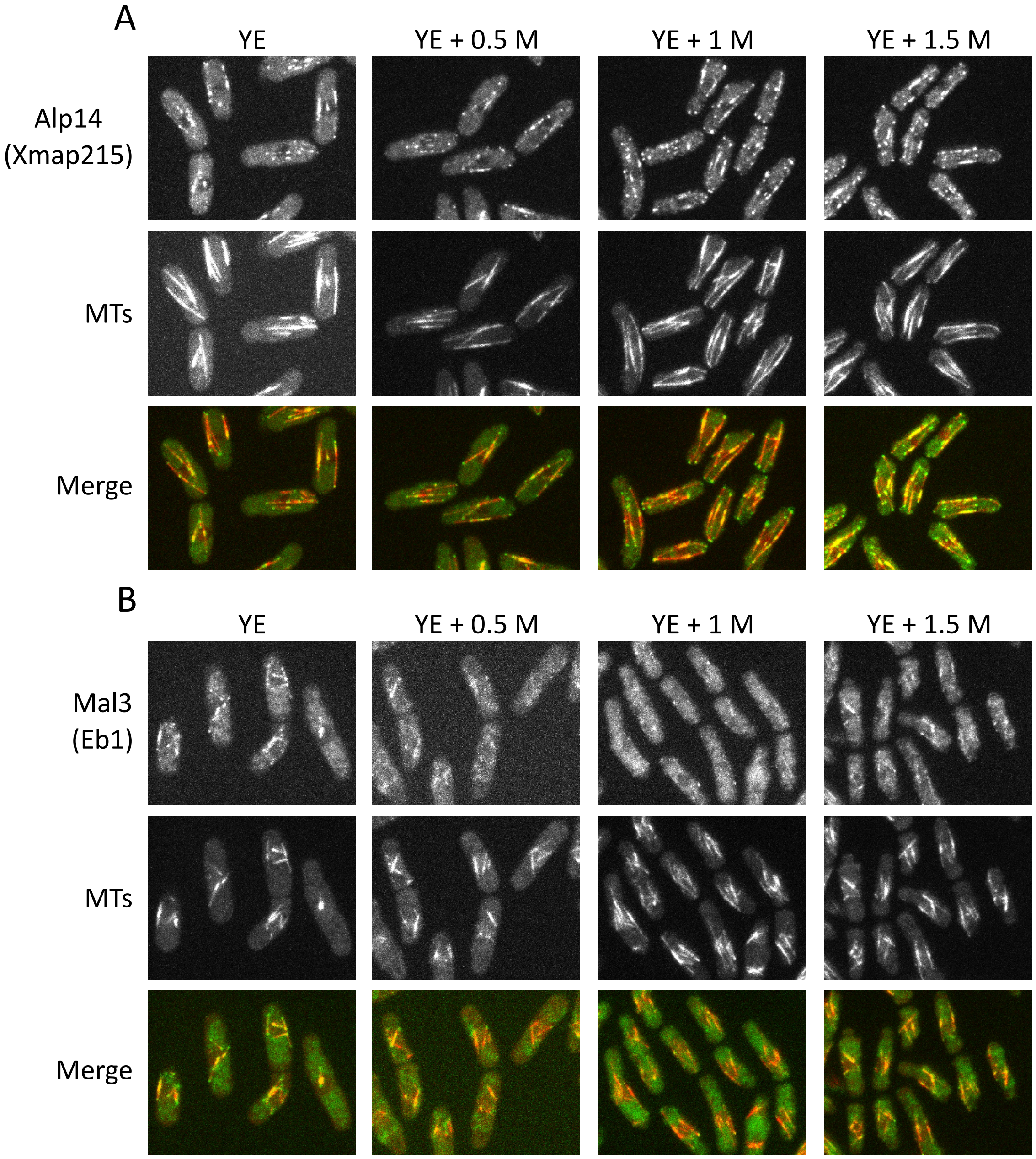
A
YE
YE + 0.5 M
YE + 1 M
YE + 1.5 M
Alp14
(Xmap215)
MTs
Merge
B
YE
YE + 0.5 M
YE + 1 M
YE + 1.5 M
Mal3
(Eb1)
MTs
Merge

Sup. Fig. 9

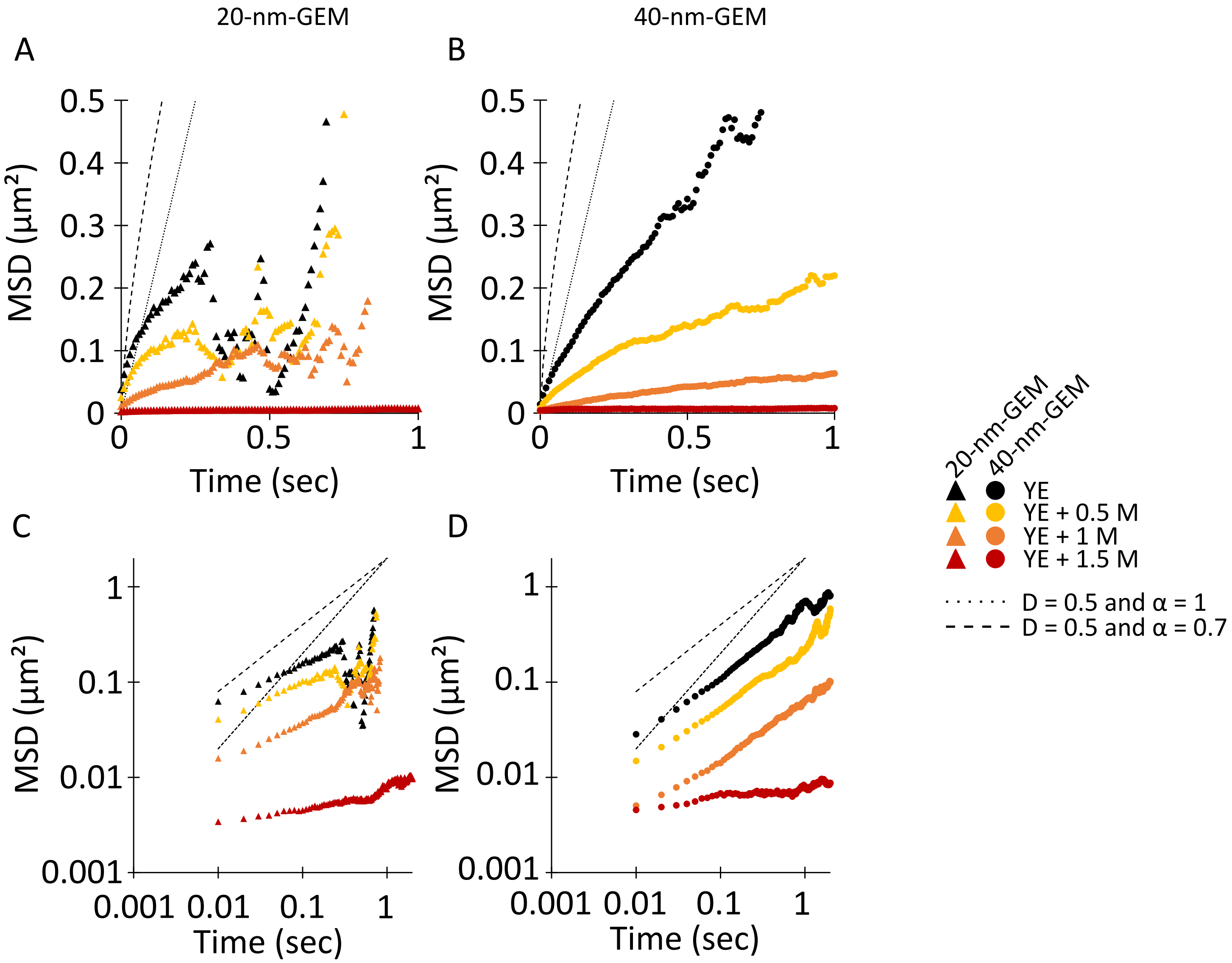

20-nm-GEM
40-nm-GEM
A
B
C
D
MSD (μm²)
Time (sec)
0.5
0.4
0.3
0.2
0.1
0
0.5
1
0.001
0.01
0.1
1
20-nm-GEM
40-nm-GEM
YE
YE + 0.5 M
YE + 1 M
YE + 1.5 M
D = 0.5 and α = 1
D = 0.5 and α = 0.7

### Sup. Table 1

| Strain | Source | Identifier |
|---|---|---|
| ***S. pombe*** | | |
| h- wt 972 | Chang lab collection | FC15 |
| h+ GFP-atb2:kanMX ade6- leu1-32 ura4-D18 | Chang lab collection | FC2861 |
| h+ sty1::ura4+ GFP-atb2:kanMX | this study | AM23 |
| h- mal3::natMX leu1-32::SV40-GFP-atb2[LEU1+] ade6- leu1- ura4- | Chang lab collection | FC2343 |
| h+ alp14::NatMX leu1-32::SV40-GFP-atb2[LEU1+] leu1- ura4- | Chang lab collection | FC2357 |
| h-rps802-GFP::kanR leu1-32 ura4-D18 ade6-216 | Knapp et al., 2019 | FC3208 |
| alp14-GFP-kan atb2-mCh::hph | this study | AM53 |
| mal3-GFP:kanMX atb2-mCh::hph | this study | AM57 |
| pREP41X_AqLs:Sapphire (20-nm-GEM) | this study | AM35 |
| pREP41X_PfV:Sapphire (40-nm-GEM) | this study | AM38 |
| h- ade6:mCherry-psy1 ish1-GFP:kanMX gpd1::hphMX6 ura4-D18 ade6 | this study | JL30 |
| h- pREp41X-Pfv-Sapphire ade6-M216 leu1-32 ura4-D18 his3-D1 | this study | JL110 |
| ***Physcomitrella patens*** | **Source** | **Identifier** |
| EB1-Citrine mCherry-tubulin | Kosetsu et al., 2013 | GPH0379#2 |

**Reagents**

| Name | Provider | Product ID |
|---|---|---|
| YES 225 | Sunrise Science Products | 2011-500 |
| Edinburgh Minimal Media (EMM) | MP | 4110-032 |
| Agar | Difco | 281210 |
| D-sorbitol D-glucose | Sigma | SLBS6381 |
| Adenine | Sigma | 1001443745 |
| Leucine | Sigma | L8000 |
| Histidine | Sigma | H8000 |
| Uracil | Sigma | U0750 |
| Carbendazim | Sigma | 378674-100G |
| Lectin | Sigma | L1395 |

**PCR primers**

| Name | Sequence |
|---|---|
| AqLs-Sapphire-F | 5’ tatagtcgctttgttaaatcatacctcgagATGCAAATATACGAAGGCAAG 3’ |
| PfV-Sapphire-F | 5’ tatagtcgctttgttaaatcatacctcgagATGCTCTCAATAAATCCAAC 3’ |
| Sapphire_R | 5’ agacattccttttacccggggatcctcgagTTATTTGTACAATTCATCAATACC 3 |
| gpd1-hph-F | 5 'GCTATGGCGTATGGGCTCATACATAAATACTTAGCTTCCCGCTCTTCTCTTTTACCAACACCGTTGAAGTgacatggaggcccagaatac 3' |
| gpd1-hph-R | 5 ' ATCATACAACATGAAACGTAGACTATGTGACCAAATAAATAAATGATAACAAGAGACTCACAAAGCACATcagtatagcgaccagcattc 3' |
